# Spatially convergent fMRI signatures of diabetes and male sex identify genetic vulnerabilities to accelerated brain aging

**DOI:** 10.1101/2025.04.14.648801

**Authors:** Anthony G. Chesebro, Luna R. Rohm, Botond B. Antal, Corey Weistuch, Helmut H. Strey, Eva-Maria Ratai, David T. Jones, Lilianne R. Mujica-Parodi

## Abstract

Age-related cognitive decline results from complex interactions between neuroendocrine and neurometabolic processes that undergo lifelong degredation, yet the mechanisms underlying these interactions remain poorly understood. This study examined the effects of diabetes and sex on functional brain networks across aging through analysis of two large cohorts (N=1,621 total) using both 3T and 7T functional MRI, complemented by spatial transcriptomic data from over 14,000 genes from six post-mortem brains. Four networks— cingulo-opercular, default mode, salience, and lateral somatomotor—exhibited significant functional decline in both individuals with diabetes and independently in males. Gene expression analysis of vulnerable networks revealed significant overexpression of insulin-dependent glucose transporters, dopaminergic and GABAergic synaptic genes, and VEGFA-VEGFR2 pathway components. These findings suggest functionally-specific circuit vulnerability to metabolic and hormonal dysregulation, potentially offering targets for early intervention before irreversible neurodegeneration occurs.

## Introduction

The maintenance of brain homeostasis in the face of extreme variability experienced over several decades of life requires neuroendocrine and neurometabolic coordination (*1–3*). Due to this inherently multi-system feedback coordination, age-related cognitive impairment and dementia are often symptomatic results of a diverse constellation of underlying pathological processes that often co-occur in ways that challenge diagnosis and prevention (*4, 5*). Final assessments of disease burden rely on measures of neurodegeneration, often informed by grey (*6*) and white matter (*7*) structural abnormalities visible with MRI, or by other imaging biomarkers of corresponding pathologies, such as tau-PET. These changes are typically diagnostic at the end stage of disease when functional decline is already clinically apparent (*4*). In the context of dementia and Alzheimer’s disease (AD), functional decline is observed much earlier on either functional MRI (fMRI) or metabolic imaging (FDG-PET) than most other biomarkers, often co-occurring with early biomarkers of disease accumulation (*8, 9*). Examining the effects of neurometabolic and neuroendocrine conditions on functional metrics is therefore an appealing prospect, as functional changes observed in aging often are sensitive to earlier disease processes and can provide insights before cognitive and behavioral impairment becomes apparent (*9*).

Disrupted energy homeostasis is a commonly implicated mechanism within the described constellation of pathological processes (*2, 10–13*). The brain maintains a delicate balance between the dominance of interregional connections for preserving functional circuits and the metabolic costs associated with maintaining these circuits (*10, 14, 15*). The brain is therefore uniquely vulnerable to disorders that impact energy homeostasis (*1, 13, 16*). Diabetes, as both an endocrine and metabolic disorder, has been increasingly implicated as detrimental in brain aging, with AD being termed in some works as “Type 3 diabetes” as the effects of disordered metabolic processes feature so significantly in age-related cognitive impairment (*10, 17–19*). Type 2 diabetes mellitus (T2DM), in particular, has been robustly associated with region-specific neurological insult and subsequent cognitive impairment (*10, 20*). As the global burden of diabetes continues to grow (*21*), understanding and mitigating the effects of this chronic degenerative disease on brain health is a high priority for public health.

Sex hormones over the lifespan also lead to striking differences between cognitive aging in males versus females (*22, 23*), often in ways that interact with other neuroendocrine conditions like diabetes (*19*). There is some evidence of an estrogen-protective effect on cognitive aging (*24*), with longer exposure to estrogen associated with preserved cognitive function in females (*25, 26*), albeit with a potentially steeper eventual decline in late age (*22*). This functionally protective effect occurs even though females typically have higher degrees of pathology associated with AD and vascular dementia (*27*). This suggests that the relative functional benefits experienced by females compared to males during aging may not necessarily be due to mitigation of underlying pathology but to different functional alterations in the face of these pathologies, such as different compensatory vascular (*28*) and metabolic (*29*) mechanisms.

Here we aim to identify the relationship between these effects by examining two large cohorts (over 1600 individuals in total) that received extensive clinical and functional MRI (fMRI) evaluations at both high (3T) and ultra-high (7T) fields using the analysis outlined in Fig. 1. First, we demonstrate that functional network segregation, known to be associated with both age- and metabolic-related changes in the brain (*15, 30*), exhibited accelerated decline in specific regions in both participants with diabetes and in males. We demonstrate that these effects are spatially congruent but are in fact independent effects not driven by baseline differences in metabolism between males and females. We also demonstrate that these effects are spatially and statistically independent from APOE4 status. Taking the regions identified as vulnerable to accelerated functional decline in both diabetes and males, we then use spatial transcriptomic data from over 14,000 genes from a post-mortem dataset (*31*) to identify candidate genes and their related pathways that could give rise to these co-localized deficits.

**Figure 1:**
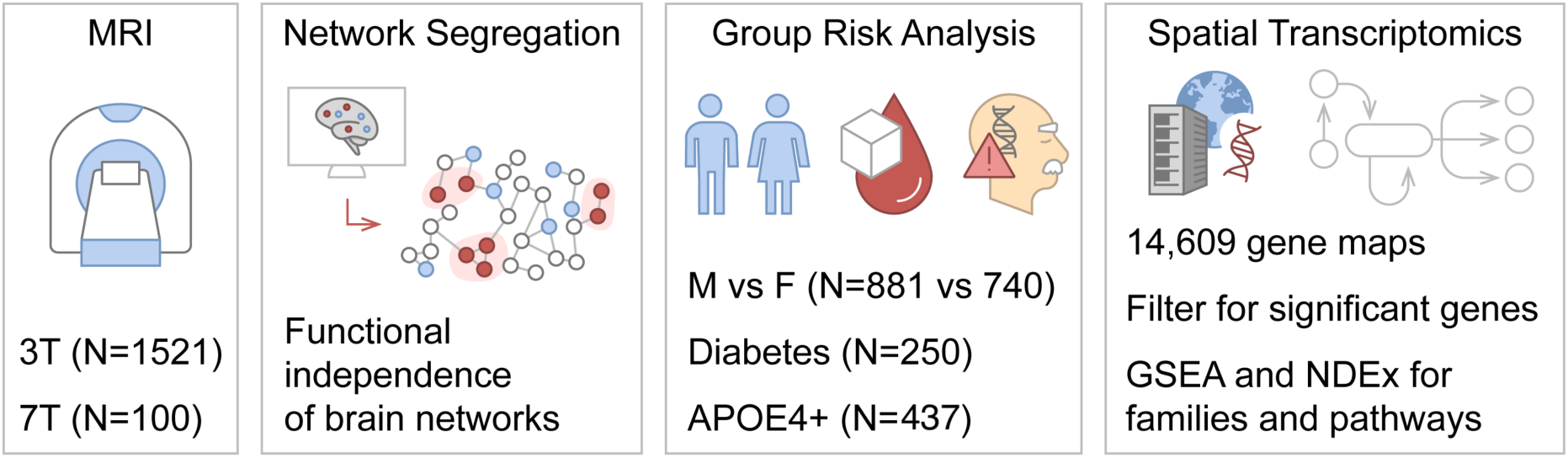
Schematic of study workflow. Two cohorts received resting-state functional MRI scans at 3T (single day) or 7T (two days, repeated scans). Both cohorts were processed to extract functional network segregation values, as this is a biomarker of age-related functional decline. We then analyzed these groups to identify spatially-localized deficits in three subgroups: males compared to females, participants with and without T2DM, and APOE4+ compared to APOE4-participants. Finally, to provide mechanistic insight to complement the functional findings, we used the AHBA (*31*) to identify genes and the associated functional pathways that are relatively over- and under-expressed in the networks that showed co-susceptibility to diabetes and male sex-related functional decline.

## Results

### Cohorts

The current study consisted of two imaging cohorts: a large cohort examined with 3T MRI and a complementary cohort scanned at ultrahigh-field 7T MRI. The 3T cohort consisted of a study of 1521 adults (829 male) aged 31-90 years old who received metabolic panels, cognitive assessment, *APOE* genotyping, and extensive diabetes monitoring. The cohort was enriched for individuals carrying the APOE4 gene (437 APOE4+) and who had well-controlled diabetes (250 diabetic, 5.6% mean HbA1c). The cohort was balanced across sexes for APOE4+ status and age. Males were slightly more likely to be diabetic (19% vs 13%) and cognitively impaired (12% vs 8%) than females, although they had equivalent HbA1c values. To ensure that the sex effects noted in this dataset were not driven by small baseline variations in diabetes prevalence across sexes, a second dataset of 100 individuals (52 male) scanned on 7T fMRI was collected. To disambiguate the effects of sex from diabetes, this cohort was recruited to be healthy at baseline (diagnosis of diabetes was exclusionary) and cognitively unimpaired. Additionally, all individuals in the 7T cohort were scanned on two different days, allowing for the reproducibility of metrics to be assessed within individuals. Complete descriptive statistics of cohort characteristics can be found in Table 1. Both cohorts received resting-state fMRI scans, as detailed in Methods.

**Table 1:** Descriptive statistics of both cohorts.

|  | 3T Dataset (N=1521) |  |  | 7T Dataset (N=100) |  |  |
| --- | --- | --- | --- | --- | --- | --- |
|  | Male | Female | Statistic | Male | Female | Statistic |
| **Sex, N** | 829 | 692 | - | 52 | 48 | - |
| **Age, mean (SD)** | 70 (12) | 69 (12) | t=-2.0,<br>p=0.04 | 45 (17) | 45 (16) | t=0.20,<br>p=0.84 |
| **Diabetes (T1 or T2), N (%)** | 160 (19) | 90 (13) | $\chi^2=11$,<br>p<0.01 | 0 (0) | 0 (0) | - |
| **Fasting blood glucose, mean (SD)** | 102 (29) | 97 (22) | t=-2.6,<br>p<0.01 | 85 (8.5) | 82 (8.5) | t=-1.8,<br>p=0.08 |
| **HbA1c** | 5.67<br>(0.78) | 5.59<br>(0.66) | t=-1.2,<br>p=0.22 | 5.17<br>(0.29) | 5.17<br>(0.33) | t=-0.07,<br>p=0.94 |
| **Cognitive impairment, N (%)** | 101 (12) | 56 (8) | $\chi^2=6.8$,<br>p<0.01 | 0 (0) | 0 (0) | - |
| **APOE4 positive, N (%)** | 223 (27) | 214 (31) | $\chi^2=3.0$,<br>p=0.08 | - | - | - |

### Age decreases functional network segregation

Functional network segregation was significantly decreased with age in both the 3T and 7T cohorts (Fig. 2). Network segregation was computed in 13 functionally defined networks (*32*) in both resting-state fMRI datasets. All network regions of interest (ROIs) are illustrated in Fig. S1. Increased age robustly correlated with decreased functional connectivity in both the 3T (Fig. 2A) and 7T (Fig. 2B) datasets. In the networks of interest for this paper (cingulo-opercular, default mode, salience, and lateral somatomotor) there are notable decreases in both datasets as age increases (r=[−0.26, −0.41] at 3T, r=[−0.27, 0.40] at 7T, p<1e-6 throughout). We report the range of correlations as the [minimum, maximum] values across networks, with the full set of values in the figures (Fig. 2 in this case) or supplemental tables (Table S1) as noted throughout the results.

**Figure 2:**
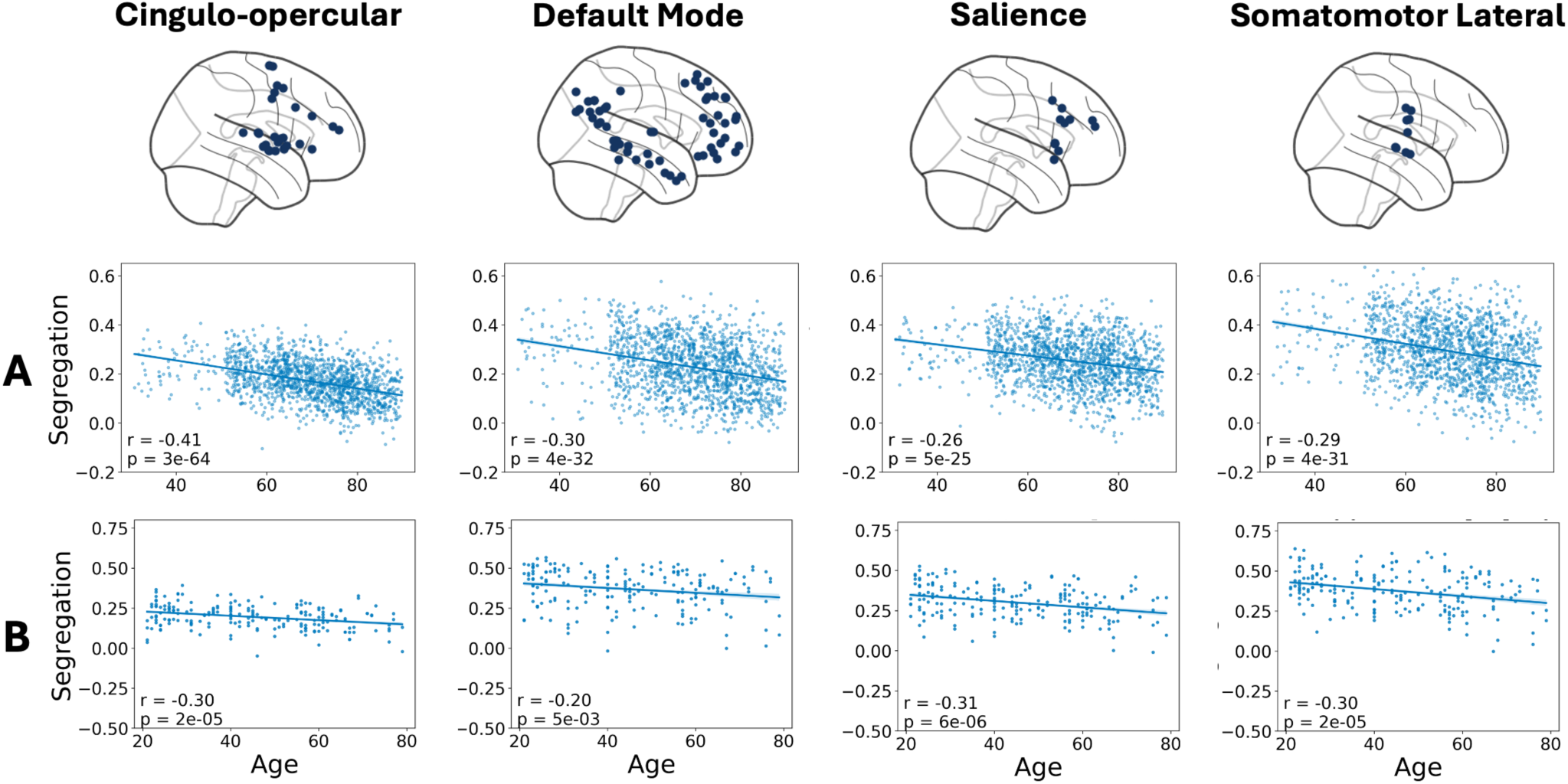
Loss of functional network segregation is associated with age throughout the brain. The first row shows the ROI locations for the functional networks of interest (cingulo-opercular, default mode, salience, and somatomotor lateral networks). Full 3D locations to visualize these coordinates are presented in Figure S1. All regression lines are presented with confidence intervals +/- standard error of the estimate. **A**. 3T fMRI dataset results show network segregations show a strong decrease correlated with increasing age. **B**. 7T fMRI dataset results show similarly strong associations of decreased network segregation with age.

In the 3T dataset, we observed this association of decreased network segregation with increased age throughout the brain (Fig. S2), with most networks showing a prominent negative correlation (r=[−0.17, −0.41], p<1e-16 throughout). Notable exceptions to this trend were reward (r=0.03, p=0.3) and dorsal somatomotor (r=0.04, p=0.4) networks; both have additional notes on preprocessing in Supplemental Methods. We also observe the association of decreased network segregation with increased age throughout the brain in the 7T dataset (Fig. S3). All networks showed a significant negative correlation with age (r=[−0.23, −0.48], p<0.01 throughout), although parieto-medial, medial temporal, and dorsal somatomotor networks showed similar non-significant results as noted in the 3T dataset and Supplemental Methods. Acquiring two scans on different days in each individual in this dataset allowed us to further test the reproducibility of network segregation and the robustness of age-related effects. Network segregation was highly reproducible within individuals across scanning days (Fig. S4; r=[0.58, 0.69], p<0.001 for all networks of interest). Additionally, correcting age-related trends with random effect of individual repeated scans did not materially change the original results (Table S1). Taken together, the 3T and 7T datasets show a robust and consistent negative association of age with network segregation throughout the brain.

### Males and participants with diabetes show decreased functional network segregation in overlapping spatial distributions

We next examined the relationship between diabetes status and network segregation across the lifespan (Fig. 3A). The 3T dataset included 250 individuals with diabetes, typically well-managed (see Table 1 and Supplemental Methods for clinical characterization). Consistent with results shown in the previous section, age generally correlated with a decrease in network segregation in participants with and without diabetes. However, participants with diabetes showed significantly lower network segregation than healthy participants in only a subset of networks, namely the cingulo-opercular, default mode, fronto-parietal, salience, and lateral somatomotor networks (t = [−2.47, 5.18], p<=0.01 throughout). These networks showed a spatial susceptibility to loss of network segregation in diabetes beyond aging; other networks did not show similar functional decreases in diabetes (Fig. S5). Additionally, these effects were not a simple correlation of diabetes status with age, as they persisted in ordinary least squares (OLS) models fit to account for age and diabetes simultaneously (Table S2).

**Figure 3:**
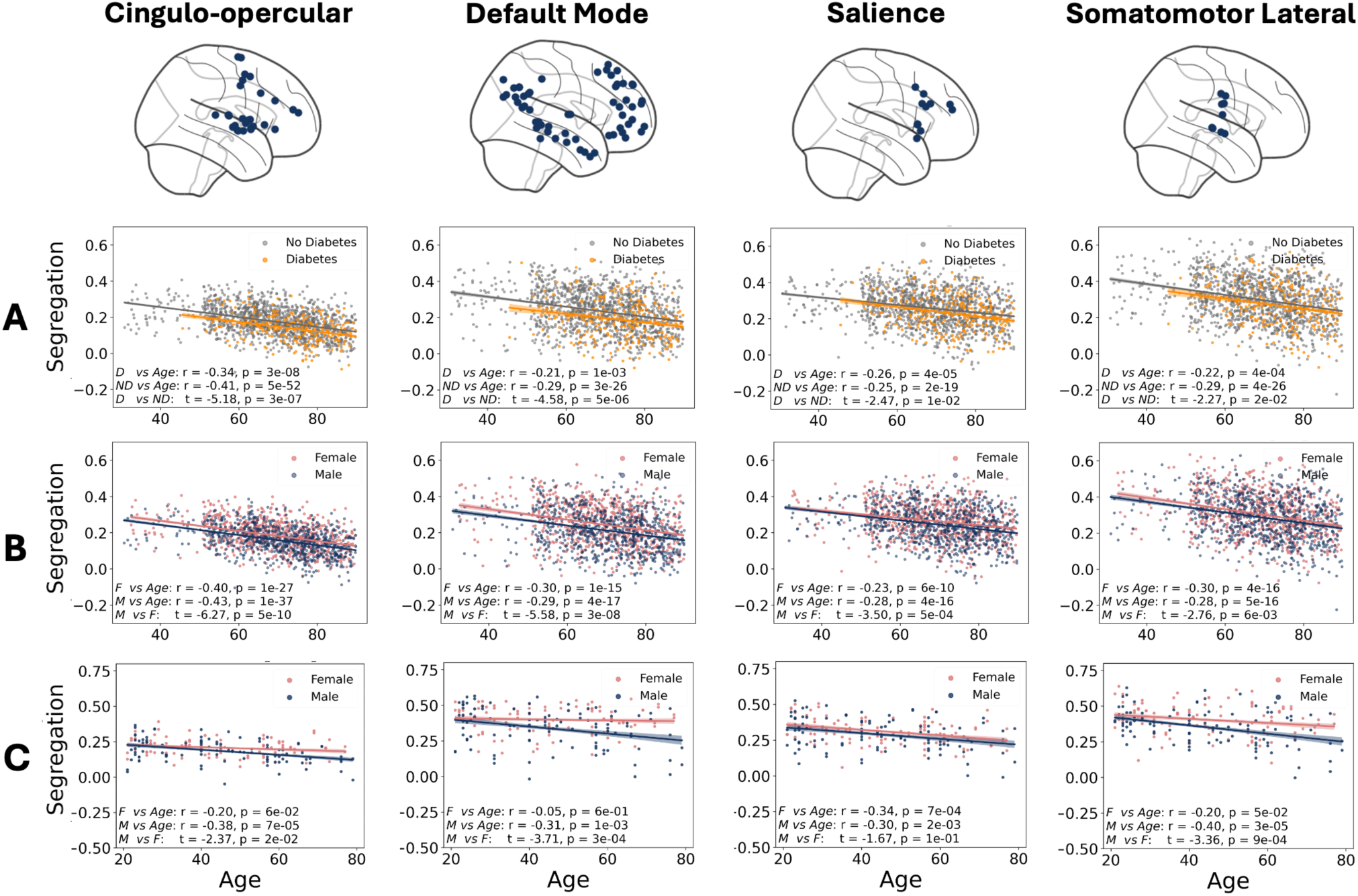
Spatial overlap between effects of diabetes and male sex on accelerated decrease of network segregation with age. Four specific networks show decreased network segregation in diabetes compared to controls as well as in males compared to females in both datasets. All regression lines are presented with confidence intervals +/- standard error of the estimate. **A**. Diabetes significantly decreases network segregation at all age points in the illustrated networks. **B**. Male sex is associated with decreased network segregation in the same networks at all ages in the 3T dataset. **C**. Male sex is associated with decreased network segregation in the 7T replication dataset as well.

Similar to diabetes, males showed a greater decrease in network segregation in a spatially dependent manner compared to females. In the 3T dataset, male sex correlated with significantly decreased network segregation across the adult lifespan in the cingulo-opercular (t=-6.27, p<0.001), default mode (t=-5.58, p<0.001), salience (t=-3.50, p<0.001), and lateral somatomotor (t=-2.76, p<0.001) networks (Fig. 3B), as well as marginally increased network segregation in the auditory network (t=1.98, p=0.05). All other networks exhibited no sex effects (Fig. S6). As with the diabetes analysis, we fit OLS models with sex and age simultaneously to ensure that sex effects were not confounded by age (Table S3). To ensure that these effects were not driven by the prevalence of diabetes across sexes, we fit an OLS with main effects of both diabetes and sex and an interaction effect between them for each network (Table S4). We observed independent main effects of sex and diabetes with no significant interaction effects, giving confidence that these effects are independent but spatially overlapping effects.

To further ensure these sex effects were independent of diabetes, we repeated the same analysis in the 7T cohort, which screened individuals to be free of diabetes at baseline. Each individual in this cohort received two resting-state fMRI scans on different days to further ensure the reproducibility of these results. Four networks showed the same spatial distribution of male sex correlated with decreased network segregation (Fig. 3C): cingulo-opercular (t=-2.37, p=0.02), default mode (t=-3.71, p<0.001), salience (t=-1.67, p=0.1), and somatomotor lateral (t=3.36, p<0.001). Other networks showed no meaningful associations with sex effects (Fig. S7). As in all other analyses, we analyzed these effects with an OLS to correct for age to ensure their independence of age (Table S3).

To ensure that these sex differences are also not caused by differences in glucose dynamics below the threshold for diagnosis of diabetes, we ran two additional OLS models for each network in both datasets, where the regressor of interest was either fasting blood glucose (collected for 689 individuals in the 3T dataset and 99 individuals in the 7T dataset) or HbA1c (collected for 689 individuals in the 3T dataset and all individuals in the 7T dataset; see Supplemental methods for clinical data notes) instead of diabetes status. These models included the biomarker (fasting blood glucose or HbA1c), sex, and the interaction effect between the two to again test for any dependencies, as well as age (Table S5). We observed interaction effects between fasting blood glucose and HbA1c with male sex only in the cingulo-opercular network and only in the 3T dataset, so there is no reproducible interaction between these biomarkers and male sex that drives the overlap between diabetes and male sex effects on network segregation.

Thus, there are four functional networks that exhibited *spatially coherent but independent* effects of diabetes and male sex (Fig. 3). These are the cingulo-opercular, default mode, salience, and lateral somatomotor networks. Each of these networks exhibited greater decreases in functional segregation beyond normal aging in individuals with diabetes, as well as exhibiting similar decreases in males independent of diabetes status. Taken together, these results indicate a potential predisposition to functional effects from endocrine pathways in these particular networks – an association that is further explored in light of APOE status and spatial transcriptomics in the next sections. Before moving on, it is interesting to note that, unlike these four networks, the fronto-parietal network shows a dichotomy between diabetes and sex. Decreased network segregation independent of age correlated with diabetes in the fronto-parietal network (Fig. S5, panel 5), but not with sex in either the 3T (Fig. S6, panel 5) or 7T (Fig. S7, panel 5) datasets.

### APOE4+ individuals showed decreased functional network segregation in different regions than diabetes and male sex effects

Given previous work establishing that APOE4 is associated with a greater risk of AD in females (*33, 34*), but potentially greater tau deposition and larger protective effects of APOE2 in males (*35, 36*), we compared the effects of APOE4 positivity (single or double APOE ε4 allele) on network segregation effects in a similar manner to the diabetes and sex effects. We observed no broad association of APOE4 status with decreased network segregation throughout the brain (Fig. S8), even when corrected for age as in the other analyses (Table S6). Two networks exhibited greater functional decline in APOE4+ individuals (Fig. 4). We observed a modest decline in parieto-medial network segregation (r=-2.15, p=0.03) and a marginal decline in the auditory network (r=01.94, p=0.05). It is interesting to note that the networks affected in APOE4+ individuals are spatially distinct from those identified as effected by diabetes and sex. While they undoubtedly lead to convergent effects in later-stage cognitive decline, the effects are spatially non-overlapping at the relatively early markers of functional decline studied in these datasets.

**Figure 4:**
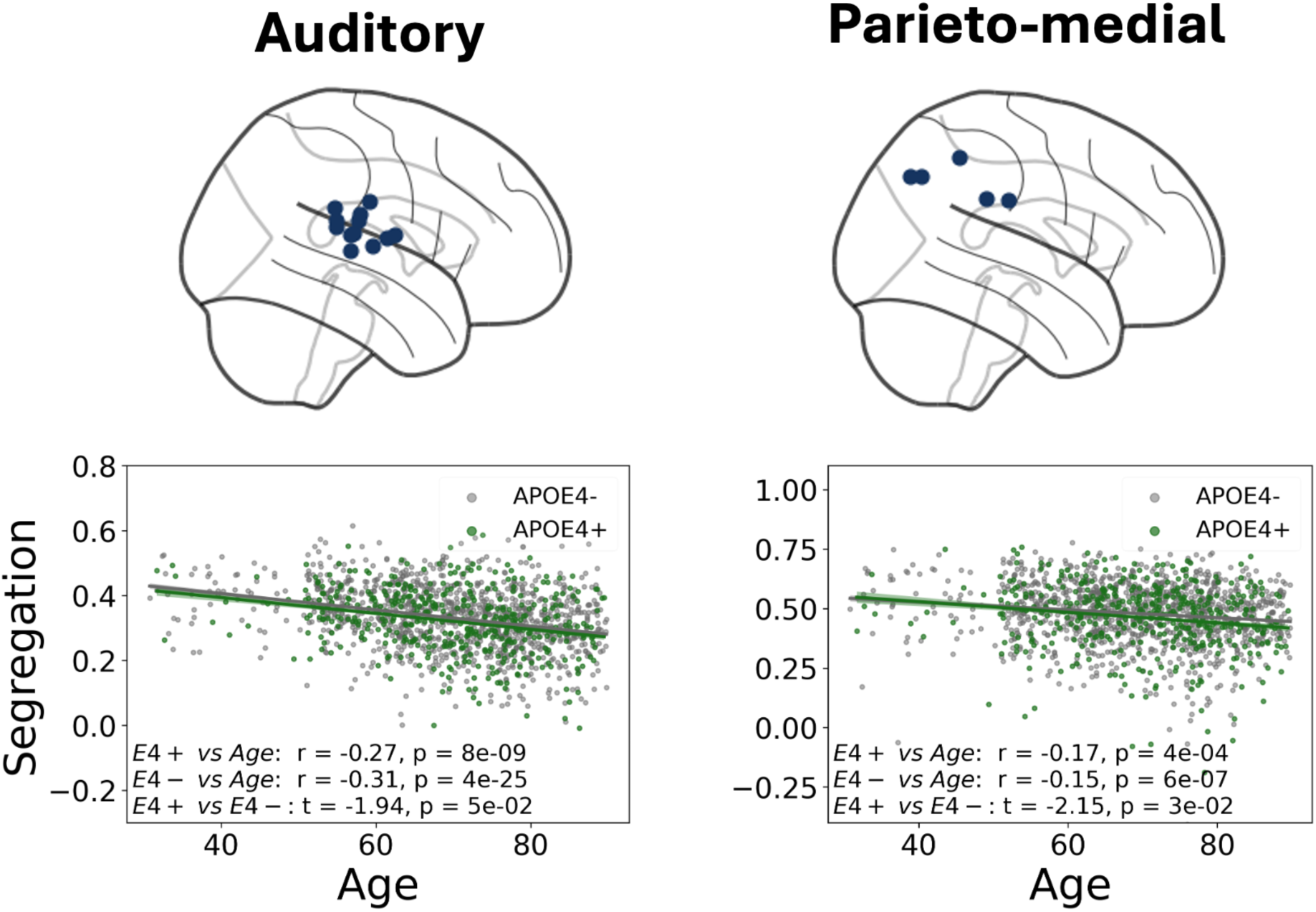
APOE4 positive status is associated with auditory and parieto-medial network functional segregation decrease. This is a notably spatially distinct pattern from the diabetes and sex effects, suggesting that they are independent pathways in early functional decline. All regression lines are presented with confidence intervals +/- standard error of the estimate.

### Targeted and unsupervised spatial transcriptomic analyses identify pathways associated with diabetes and male sex network segregation effects

We used spatial transcriptomics to extend these results and explore what genetic pathways are expressed more in the networks that exhibited functional decline in these conditions. We conducted these analyses on transcriptomic data from six post-mortem brains provided by the Allen Human Brain Atlas (*31*). The measure of interest was the relative expression ratio of networks of interest (cingulo-opercular, default mode, salience, and lateral somatomotor) compared to the rest of the functionally-defined ROIs. We used this ratio to determine which genes were most significantly associated with the functional decline in diabetes and male sex. A relative expression level of 1 indicated the same level of expression (no association with networks of interest), while an expression significantly greater or less than 1 indicated relative over- or under-expression, respectively, of a gene in these networks compared to the rest of the brain. These analyses followed two analysis streams: a targeted set of genes selected on *a priori* hypotheses and an unsupervised approach based on analyzing relative expression levels of all available genes (14,609 genes in total).

In the targeted analysis, we tested three initial hypotheses: the overexpression of insulin-sensitive glucose transporters in regions affected by diabetes, the overexpression of estrogen receptors in regions adversely affected by male sex, and the overexpression of metabotropic glutamate receptors (mGluR) in regions adversely affected by male sex. We hypothesized genes encoding insulin-sensitive glucose transporters (*SLC2A4, SLC2A4RG, SLC2A12*) would be overexpressed in the regions that experienced functional decline in diabetes, while other glucose transporters (e.g., *SLC2A1, SLC2A3*) that are broadly expressed in neurons would have an unrelated spatial distribution, as prior work has found that insulin-dependent glucose transport, particularly via GLUT4, supports functional neurometabolic demands (*37, 38*). This hypothesis was borne out by the spatial transcriptomic data (Fig. 5A, upper panels, and Fig. 5B), with insulin-sensitive glucose transporters overexpressed (ratio=1.37, [1.27, 1.51]) compared to other glucose transporters (ratio=1.11, [0.92, 1.28]) in these networks. As estrogen presence, either through normal hormonal variation or hormone replacement therapy, has been associated with improved cognitive outcomes in aging (*24–26*), we hypothesized that the genes encoding estrogen receptors (*ER1, GPER1*) would exhibit increased expression in the regions that showed a female protective effect when compared to selected control hormone receptors (inhibin receptors; *INHA, INHBA, INHBB, INHBC*) known to be widely expressed throughout the brain (*39*). While we did not find the control genes to be meaningfully associated with the selected networks (ratio=1.02, [0.85, 1.29]), we also found the estrogen receptors to not be significantly increased in the networks of interest (ratio=1.02, [1.01, 1.04]). We tested a final targeted hypothesis on mGluR overexpression in the regions exhibiting a potential female protective effect, as estrogen is known to modulate the activity of mGluRs (*40–42*). Broadly, we observed genes encoding for mGluRs (*GRM1-GRM8*) to be overexpressed in the regions exhibiting a female protective effect (ratio=1.16, [1.01, 1.40]), especially when contrasted with serotonergic receptors that are expressed throughout the brain (ratio=1.05, [0.85, 1.29]). As a summary of the targeted analysis results: while insulin-dependent glucose transport and mGluR expression levels were relatively overexpressed in the regions shown to be adversely affected by diabetes and male sex, the relative expression levels of estrogen receptors did not have a reliable association with these regions, indicating a functional role of hormones other than simple presence or absence.

**Figure 5:**
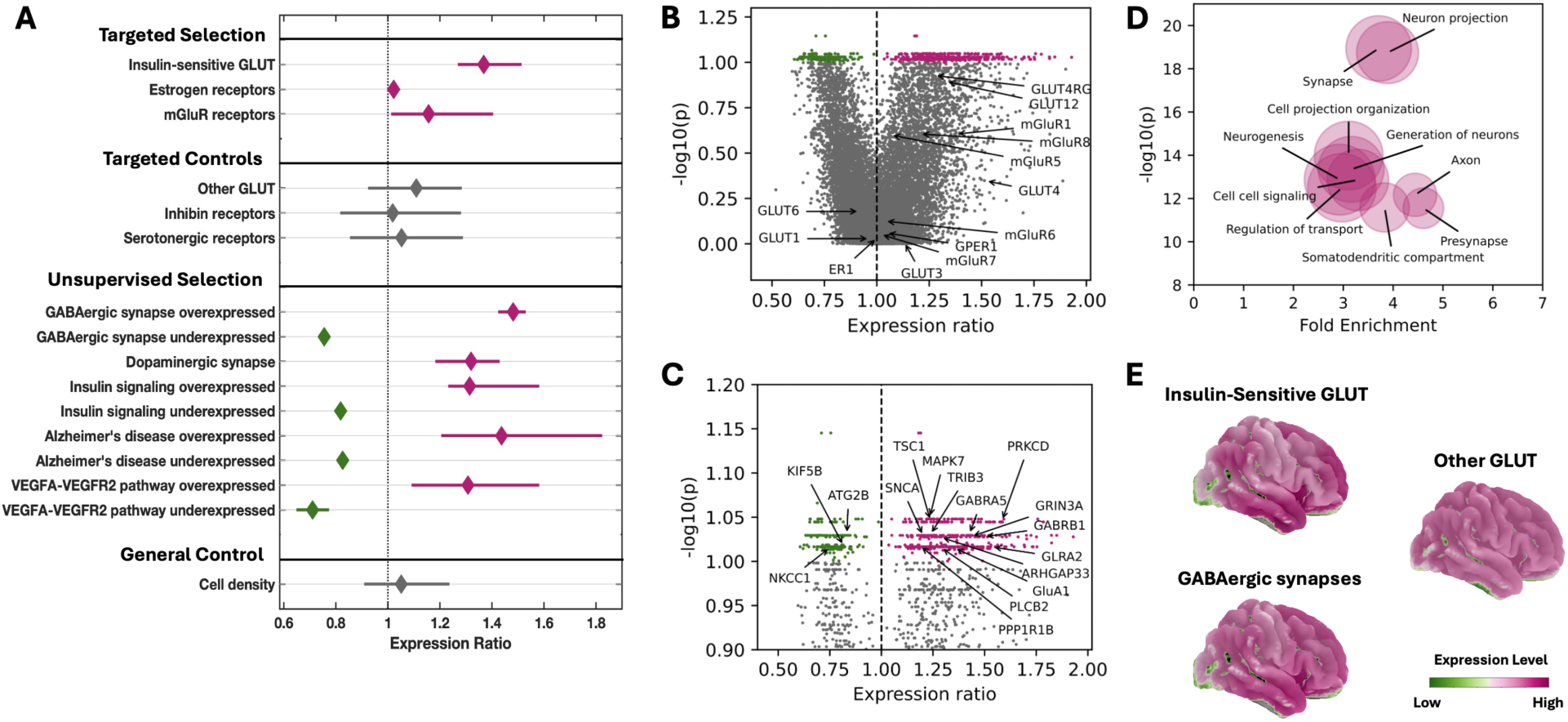
Spatial transcriptomics implicate insulin signaling, synaptic activity, and vascular signaling in region-specific functional decline. **A**. Relative expression levels of the different genes selected for targeted hypotheses and controls, as well as those shown to have significantly higher expression by random selection. Full gene lists can be found in Table S7. Expression ratio is the relative level of mRNA expression in the four networks of interest compared to the rest of the brain. Bars are drawn showing the range of expression levels, with diamond centers at the mean expression level. **B**. Entire set of relative expression levels for all genes, with p-values computed based on spatially-correlated null surrogate maps (*81*). Specific genes from the targeted hypotheses are shown with arrows, with selected genes typically having expression ratios well above 1, while control genes are relatively uniform throughout the brain. **C**. Zoomed section of the upper portion of B, with specific genes from the unsupervised selection highlighted. **D**. GSEA results showing the top 10 associated gene set collections in the full, unsupervised analysis, implicating a predominance of neuronal function and extracellular signaling pathways. **E.** Example spatial distributions of specific gene expression levels to illustrate the predominantly temporal, with muted frontal and medial parietal, locations of the relative overexpression of genes associated with the diabetes and sex effect networks. The genes are selected to highlight insulin-dependent glucose transport (top left) and synaptic activity (bottom left) that occur in spatially overlapping patterns. By contrast, genes encoding for GLUT1 and GLUT3, present throughout virtually all neurons, show no meaningful regional variability in expression.

To complement these targeted analyses, we performed an unsupervised analysis across the entire set of genes available through the AHBA to identify the pathways most significantly associated with the regions identified as susceptible to functional decline in diabetes and males. The expression ratio and relative significance level (with p-values adjusted via FDR-correction) were computed independently for every gene available (Fig. 5B shows the entire volcano plot of genes with the targeted selection genes highlighted, while Fig. 5C shows a zoomed portion with genes highlighted by the unsupervised analysis as most significant). To aid in interpreting the genes most significantly associated with the networks affected by diabetes and male sex, we performed two additional analyses on the top 500 genes (selected by significance level): gene set enrichment analysis (GSEA; (*43*)) and pathway search via Network Data Exchange (NDEx; (*44*)). NDEx results highlighted genes in insulin signaling pathways in these regions (*ARHGAP33, MAPK7, PRKCD, TRIB3, TSC1*) and general pathways associated with AD (*PLCB2, SNCA, FZD1*). Interestingly, one gene was common between the insulin-signaling and AD-related pathways (*KIF5B*) and was relatively under-expressed in these networks. NDEx also highlighted synaptic signaling pathways associated with these networks, with GABAergic (both relative over- and under-expression of different genes) and dopaminergic (relative over-expression only) synapses in particular, and no serotonergic receptor genes noted (signifying they are valid control genes). Finally, the NDEx pathways showed a number of genes in the VEGFA-VEGFR2 pathway as associated with the spatial distribution of diabetes and sex effects noted above, with seven genes significantly over-expressed and two under-expressed in these regions. The spread of expression ratios of the genes highlighted by the NDEx analyses is shown in the bottom of Fig. 5A, with relative expression ratios and FDR-corrected significance levels shown in the zoomed volcano plot in Fig. 5C. We provide a complete list of all genes highlighted by these analyses with the pathways they belong to in Table S7. GSEA analysis revealed a similar pattern of related gene families (Fig. 5D), implicating a predominance of distal association (presynapse, synapse, somatodentritic compartment, neuron projection gene sets) as well as intercellular communication (cell-cell signaling gene set) in the pathways most associated with overexpression in these regions. The entire list of gene families as found by GSEA is provided in Table S8.

The spatial distributions of expression of the insulin-dependent genes and GABAergic synapses illustrate how these processes overlap spatially independently of cell density (Fig. 5E). The spatially localized distributions of both diabetes-associated genes (*SLC2A4, SLC2A4RG*) and genes associated with GABAergic synapses (*GABRA5, GABRB1, GLRA2*) contrasted with control genes for glucose transporters (*GLUT1, GLUT3*) that are expressed broadly throughout the brain. Both the targeted and unsupervised gene selections showed a temporal predominance in expression, with medial frontal and parietal highlights as well. In contrast, the control visualization of insulin-independent glucose transporters showed a relatively uniform distribution throughout the brain. As a final control analysis for both the targeted and unsupervised analyses, we computed the relative expression levels of genes that are considered markers of cell density throughout the brain to demonstrate that the effects seen here are not driven by cell density (Fig. 5A, bottom panel; ratio=0.95, [0.91, 1.24]). Discussion of the rationale for the selection of these control genes is provided in Supplemental Methods, and all expression ratios with associated p-values (both corrected and uncorrected) for the entire set of genes analyzed can be found in Table S9.

In summary, the regions that exhibit spatially overlapping effects of diabetes and male sex (Fig. 3) showed a relative overexpression of genes associated with both insulin signaling and inter-neuron communication, especially dopaminergic and GABAergic signaling (Fig. 5). This is in contrast to no noticeable differential expression of estrogen receptors, indicating that although these regions exhibited a protective effect in females, this was not driven by baseline estrogen receptor density. The involvement of cell-cell signaling pathways, both endocrine and neuronal, further supported the hypothesis that these findings were driven by functional changes early in pathologic processes rather than end-stage structural degradation causing broad disruption. This was further supported by the finding that these regions also have baseline differences in the VEGF-VEGF2A expression levels, as vascular changes are also typically observed before structural integrity is compromised.

## Discussion

Together, our analyses suggest overlapping neuroendocrine and neurometabolic effects on functional decline during aging. First, we showed a strong replication demonstrating increased age correlated with loss of network segregation, measured at both 3T and 7T resting-state fMRI, across a combined sample of more than 1600 individuals. This is an unsurprising finding given the prior evidence for this association (*15, 30, 45–48*), but nonetheless helpful to determine the utility of network segregation in this context, as it is a broad measure of functional dependence between regions (*49*). This also complements recent findings that functional connections that are metabolically expensive (*11, 50*), especially over a longer distance (*14, 49*), are weakened with age – a relationship that is captured by network segregation (*15*). As functional changes are some of the first to appear in age-related and neurodegenerative processes like AD (*8, 9*), it is necessary to establish functional imaging biomarkers related to early pathological changes, both for basic research and for potential diagnostic purposes.

We also presented evidence of co-susceptibility to both diabetes and male sex effects on worsening network segregation beyond normal aging in the cingulo-opercular, default mode, salience, and lateral somatomotor networks. This co-localization unifies many previous threads of evidence into a cohesive picture of neuroendocrine coordination to maintain homeostasis. For example, there is prior evidence for task-specific sex differences in cingulo-opercular (*47, 51*) and salience (*47, 52*) network function, albeit often in the context of other neurological conditions (*52–54*). These effects are noted, however, without mention of metabolic dysregulation. Similarly, both type 1 diabetes (*55*) and T2DM (*10*) accelerate brain aging, particularly in subcortical regions (*10*), but with limited assessment of region-specific functional deficits, particularly not assessing the combined effects of sex and diabetes in co-localized deficits. In this context, the evidence presented here is particularly striking, as it implicates the same regions as vulnerable to diabetes and male-specific insult or female-specific protective effects, discussed further below. We noted that the fronto-parietal network exhibited decreased functional segregation in diabetes but not in males, suggesting there may be effects unique to diabetes present in this network. There is some evidence for frontal-specific effects of T2DM (*10, 56*), particularly in acute metabolic insult (*57*), but further research could elucidate why this network remains independent of male-specific functional decline as well.

One prominent potential mechanistic driver of sex-specific functional effects is the presence of the APOE4 mutation, known to drive sex-related differences in neuropathology (*58, 59*). In the analyses presented here, however, the deleterious effects of APOE4 on network segregation were spatially distinct from the regions implicated in both sex and diabetes analyses and are confined to predominantly parieto-medial with some auditory network involvement. Given the relatively weak associations of APOE4 with functional decline overall, the evidence presented here does not offer much insight into the impact of APOE4 on the interplay between sex and diabetes effects and merely reaffirms its previously reported regional effects in the temporal and parietal lobes (*60, 61*). These results do, however, remove APOE4 from consideration as a strong mechanistic driver of the effects observed in this study.

The spatial transcriptomic results we present, while not causative evidence, provide several interesting markers for future exploration of mechanistic pathways underlying neuroendocrine feedback, particularly in the context of neurometabolic dysregulation. The relative over-expression of insulin-dependent glucose transporters in regions affected by diabetes reaffirms recent *in vitro* findings that implicate these transporters, especially GLUT4, in crucial supporting roles during periods of increased neurometabolic demand (*37, 38*). The implication of insulin signaling pathways in diabetes-related networks is perhaps unsurprising, but the genes that arose during the unsupervised analysis as having spatial co-localization with the networks associated with diabetes effects provide some interesting insights. The genes *PRKCD* and *TRIB3* in particular have been implicated in neurovascular complications in diabetes (*62, 63*), but this specific spatial distribution has not previously been reported or analyzed. Similarly, lowered *KIF5B* has been associated *in vitro* with lowered dendritic function (*64*), synaptic plasticity (*65*), and membrane localization of GLUT4 (*66*). The finding of lowered baseline expression of *KIF5B* in the regions affected by diabetes underscores its importance as a cofactor in insulin signaling, with its relative under-expression observed in regions exhibiting diabetes-specific functional deficits. These unsupervised analyses also provide increased evidence for the Wnt signaling pathway (specifically the finding of increased expression of *FZD1*), already frequently implicated in aging and AD (*67*).

Although the estrogen-protective effect has been discussed in previous literature (*24*), we did not observe an over-expression of estrogen receptors in the networks that exhibit greater functional decline in males. We did, however, observe relatively higher expression of mGluRs, which are known to have functional dependence on the presence of estrogen (*41, 42*), indicating that the differences between male and female participants may be driven by functional pathways related to sex hormones without being directly tied to the hormone receptors. This was further supported by the implication of other specific neurotransmitter signaling pathways, particularly *GABRA5* over-expression, which has been associated with sexual dimorphisms in both GABAergic signaling and glucose regulation in mouse studies (*68*), even though it is equally expressed across sexes (*69*). Also associated with these regions were genes encoding dopaminergic synapse pathways, including *GRIA1* which is hypothesized to be functionally dependent on androgen hormones (*70*). These effects were not driven by simple increased neuronal or synaptic density, as both markers of neuronal density and serotonergic receptors did not differ between these networks and the rest of the brain (Fig. 5A). Finally, we noted that there are several genes from the VEGFA-VEGFR2 pathway over-expressed in these networks of interest at baseline. Although the mechanism of this angiogenesis pathway in supporting healthy brain aging is incompletely characterized, emerging evidence has shown a striking sexual dimorphism in its expression during brain aging, with females having a protective effect of increased angiogenesis up to age 75 (*28*).

In spite of the large sample sizes included in this study, there remain several limitations that should be addressed in future work, particularly if focused on personalized medicine. Network segregation, while a useful metric to assess age-related functional decline, was originally designed as a resting-state metric and is therefore limited in probing task-specific networks. It is also important to note that the datasets included here are cross-sectional by design and therefore are not as indicative of individual changes (although they are demonstrated here to be reproducible within individuals, and demonstrated previously in (*71*)). An ideal study to address both of these limitations would be a targeted set of task-based fMRI sequences to probe different circuits known to be impacted by aging and repeated in the same individuals over a longer time period to assess how these metrics can change within individuals. Finally, it should be noted that while spatial transcriptomics provide an excellent sampling of baseline distributions of mRNA expression, protein expression levels fluctuate with age in specific patterns (*72, 73*). A more thorough characterization of how these correlate with functional decline as shown here would look at fluid-based biomarkers to assess dynamic protein levels to complement the spatial transcriptomic maps.

The analyses we present here identified four key networks (cingulo-opercular, default mode, salience, and lateral somatomotor) that showed particular vulnerability to both diabetes and male sex, independent of normal aging effects. These findings were supported by spatial transcriptomic analyses showing overexpression of insulin-sensitive glucose transporters and dopaminergic and GABAergic receptors in these regions, while also highlighting the involvement of specific synaptic and vascular signaling pathways. Notably, while APOE4 status showed some influence on brain function, its effects were spatially distinct from the diabetes and sex-specific patterns observed. The overlap between diabetes and male sex effects in specific brain networks, along with the identification of relevant genetic pathways, provides new insights into the co-localized neuroendocrine and neurometabolic mechanisms of functional decline in aging and suggest potential targets for personalized interventions based on metabolic and endocrine profiles.

## Materials and Methods

### Participants and fMRI Acquisition – 3T dataset

The participant cohort for the 3T fMRI dataset is composed of data from the Mayo Clinic Study of Aging (MCSA), which has previously been described in detail (*74*). The subset of data analyzed in this study contained data for 1,521 participants between 31 and 90 years old. Complete clinical data acquired, including diabetes status, fasting blood glucose, HbA1c levels, cognitive impairment, and APOE4 status is reported in Table 1. Additional details of participant recruitment and clinical data acquisition are included in (*74*), with additional notes about the specific data included in this study in the Supplemental Methods as well. Structural and functional MRI scans in this dataset were acquired on a 3T GE Discovery MR750. The imaging protocol included an echo-planar imaging (EPI) sequence for whole-brain blood oxygen level-dependent (BOLD) signals and a T1-weighted imaging sequence for structural images. BOLD acquisition parameters comprised a repetition time (TR) of 2.9s, echo time (TE) of 30ms, flip angle of 90°, voxel size of 3.28×3.28×3.3mm, and total acquisition length of 7min 44s (160 volumes). T1-weighted structural volumes were acquired with 1.2×1.02×1.02mm voxel size, TE=3.044ms, TR=7.36 ms, and flip angle of 8.0°.

### Participants and fMRI Acquisition – 7T dataset

A second, independent fMRI dataset was used to evaluate the replicability of the observed age and sex-specific associations with functional network segregation. This dataset was acquired at 7T field strength from a cohort of N=100 healthy adults aged between 21 and 79 years. This study was conducted under the oversight of the Internal Review Board at Massachusetts General Hospital (Boston, MA, USA) and State University of New York at Stony Brook (Stony Brook, NY, USA). Informed consent was obtained prior to a physical examination and administration of cognitive testing to ensure cognitive health (see Supplemental Methods for details). Each participant was scanned on two separate occasions, at least 24 hours apart, using identical procedures.

Structural and functional MRI scans in this dataset were acquired on a 7T Siemens Magnetom (Siemens Healthineers, Erlangen, Germany). Similar to the 3T dataset, the imaging protocol EPI-BOLD fMRI and T1-weighted structural sequences. BOLD images were captured using a protocol for detecting resting-state networks, with acquisition parameters optimized via a dynamic phantom (BrainDancer; ALA Scientific Instruments) (*75*). The resulting protocol for BOLD acquisition included a simultaneous multi-slice (SMS) slice acceleration factor of 5, R=2 acceleration in the primary phase encoding direction (48 reference lines), and online generalized autocalibrating partially parallel acquisition (GRAPPA) image reconstruction. Additional acquisition parameters comprised a repetition time (TR) of 802ms, echo time (TE) of 20ms, flip angle of 33°, voxel size of 2×2×1.5mm, and total acquisition length of 10min (750 volumes). T1-weighted structural volumes were acquired with 1mm isotropic voxel size and four echoes using a protocol with TE1=1.61 ms, TE2=3.47ms, TE3=5.33ms, TE4=7.19ms, TR=2,530ms, flip angle of 7.0°, R=2 acceleration in the primary phase encoding direction (32 reference lines), and online GRAPPA image reconstruction.

### fMRI preprocessing, network segregation, and statistical methods

The fMRI data from both the 3T and 7T datasets were preprocessed using a combination of methods from *fMRIprep* (*76*), SPM (SPM12, UCL) and the *nilearn* Python library (*77*) as described in previous work (*12*). The T1-weighted anatomical images first underwent bias field correction, skull-stripping, and normalization to Montreal Neurological Institute (MNI) templates. Functional BOLD contrast images were then realigned to correct for head motion, slice-time corrected, and adjusted for geometric distortions caused by magnetic field inhomogeneities by registering the BOLD reference to the intensity-inverted T1-reference (*78*). Following this, the functional images were coregistered with the anatomical scans and normalized to MNI space. To minimize physiological noise, the time-series were bandpass filtered between 0.01 Hz and 0.1 Hz, and mean signals from white matter and cerebrospinal fluid voxels were regressed out. Additionally, six motion parameters were also regressed out reduce the effect of motion-related artifacts. Spatial smoothing was applied to the data using a Gaussian kernel with a full width at half maximum (FWHM) of 5mm. Finally, the time-series were parcellated into 300 functional regions of interest using the Seitzman atlas for subsequent analyses (*32*).

Network segregation was computed as the difference of within network and between network connectivity, given as previously defined in (*15*) by

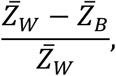

where *Z̄_W_* and *Z̄_B_* are the average Fisher-transformed correlation for nodes within and between networks, respectively. It is important to note that this average is of positive correlations only, and that all positive correlations are used in the average, regardless of weight.

Associations with age as presented in Fig. 2 are simple Pearson correlations with linear regressions and the standard error of the regression visualized as well. For all comparisons of variables of interest (diabetes, sex, APOE4 status, repeated scans) presented in Fig. 3-4, Fig. S2-S8, and Tables S1-S6, ordinary least squares (OLS) regression for multiple independent variables or random effects were employed as appropriate. All models that assessed interaction effects included main effects of both variables as well. Additional statistical methods to assess significance of spatial overlap with gene maps are discussed further below.

### Gene expression analyses

Regional microarray expression data were obtained from six post-mortem brains (one female, 24– 57 years old, 42.5±3.4) provided by the Allen Human Brain Atlas (*31*) (AHBA, https://human.brain-map.org). Data were processed with the *abagen* toolbox (version 0.1.3; https://github.com/rmarkello/abagen) using a 300-region functionally-defined volumetric atlas in MNI space (*32*). Our analysis pipeline followed the standard *abagen* pipeline with minor variations as detailed in Supplemental Methods. This process resulted in expression maps for 16,826 genes throughout the brain. These resulting expression maps were then filtered for genes included in brain-specific gene sets (*79*), selecting 87% (14,609) of the gene maps for subsequent analyses.

To determine which genes were most associated with the functional regions most affected by diabetes and male sex, the average expression levels were computed for these ROIs (cingulo-opercular, default mode, salience, and somatomotor lateral networks) and then ratioed with the expression levels for the remaining ROIs to compute the relative over/under expression of the genes in these ROIs compared with the rest of the brain. To correct for potential spatial autocorrelations within the gene maps, the toolbox *neuromaps* (*80*) was used to create null spatial maps exhibiting similar autocorrelation patterns (*81*). For each gene selected, 1,000 null maps were produced and the ratio between affected and unaffected regions were computed in the same way as the true ratios, yielding a distribution of expression ratios across all permutations. The resulting ratios were combined, and a gamma distribution was fit to them, allowing for the quantification of statistical significance between the observed gene expression ratio and the potential distribution from null maps. To address multiple comparisons, the p-values obtained from this fit were FDR-corrected (*82*). An initial analysis was done using the same method for the APOE-associated functional networks (both parieto-medial alone and parieto-medial and auditory networks combined). As this analysis yielded extremely weak associations with all genes, it is excluded from the results.

Using the ratios and statistical significance values calculated from the above steps, two analyses were then performed: a targeted analysis of genes hypothesized *a priori* to be potentially related to pathways of interest, along with control genes that should show no significant correlations with the contrasts of interest; and an unsupervised analysis consisting of taking the genes most significantly associated with the spatial patterns observed in the functional results and analyzing which pathways these belonged to. For the targeted analysis, genes hypothesized to be associated with diabetes (e.g., *SLC2A4, SLC2A12*) and sex-specific processes (e.g., *ESR1, GRM1*) were selected and compared to control genes that should have no spatial relationship with the effects of interest (e.g., insulin-insensitive glucose transporters, inhibin receptors). For the unsupervised analyses, the top 500 most statistically significant genes (both over and under-expressed in the diabetes and sex-specific effect networks) were used in a gene set enrichment analysis (GSEA; (500 genes being the maximum cutoff allowed by the tool). Standard GSEA software (https://www.gsea-msigdb.org; (*43*)) was used for this analysis, specifically focused on C5 (ontology gene sets) overlap from the molecular signatures database (MSigDB; (*83*)). This unsupervised approach was extended using the same 500 genes as a query for related pathways using the Network Data Exchange (NDEx; (*44*)).

Spatially smoothed gene maps provided in previous work (*84*) were used to visualize the spatial distributions of the genes in the targeted analysis and discovered in the unsupervised analysis contrasted with the spatial distribution of the control genes. The targeted pathway shows the insulin-sensitive glucose transporter pathway genes (*SLC2A4, SLC2A4RG*); the unsupervised pathway shows the distribution of GABAergic synapse related genes (*GABRA5, GABRB1, GLRA2, SLC12A2*); and the control glucose transporters show insulin-independent glucose transporters that are widely expressed in neurons and astrocytes (*SLC2A1, SLC2A3*). Visualizations involved normalizing the original maps to the unit range and averaging across genes, then plotting in on an average 3D surface using the *nilearn* package (*77*).

## Acknowledgments

LRR acknowledges the Berlin-Boston Research Exchange (http://www.berlinboston.com).

## Funding

Personnel support for this study included:

National Institutes of Health NIHGM MSTP Training Award T32-GM008444 (AGC).

The data contained in the 3T dataset were obtained under one of the following:

National Institutes of Health U01AG06786 (Ronald Petersen, PI).

National Institutes of Health P50 AG16574 (Ronald Petersen, PI).

## Author Contributions

Conceptualization: AGC, BBA, LRMP Methodology: AGC, BBA, CW Investigation: AGC, LRR, BBA Visualization: AGC, LRR, BBA Funding acquisition: EMR, LRMP

Project administration: HHS, DTJ, EMR, LRMP Supervision: CW, HHS, EMR, DTJ, LRMP Writing – original draft: AGC, LRR, BBA

Writing – review & editing: AGC, LRR, BBA, CW, HHS, EMR, DTJ, LRMP

## Competing interests

Authors declare they have no competing interests.

## Data and materials availability

The 3T fMRI dataset used in these analyses is part of the Mayo Clinic Study of Aging (MCSA), and the data are publicly accessible upon reasonable request to the MCSA. The 7T fMRI dataset used in these analyses is part of the Protecting the Aging Brain dataset, which is provided by the authors at openneuro.org/datasets/ds005405. All spatial transcriptomic data are provided with open access by the Allen Institute at https://human.brain-map.org. Processed spatial transcriptomic maps in MNI space and parcellated for FreeSurfer have also been provided at https://www.meduniwien.ac.at/neuroimaging/mRNA.html. All fMRI preprocessing and statistical results are done using open-source packages as described in the methods, and all results from custom analyses (e.g., relative gene expression ratios in particular networks) are included as supplemental data files.

## Supplementary materials

### Supplemental methods

#### Clinical data collection

For Table 1, not all participants in the 3T dataset received fasting blood glucose HbA1c measurements. The entire sample was 689 (372 males), with the statistics as noted in Table 1. All individuals in the 3T dataset had diabetes status confirmed by clinical diagnosis, with 244 individuals (156 male) confirmed to have Type 2 diabetes, and 6 individuals (4 male) confirmed to have Type 1 diabetes. For the 7T dataset, participants were recruited from the Boston metropolitan area through advertisements. Subject were excluded if they had contraindications to MRI, neurological or psychiatric conditions, a history of brain injury, insulin resistance, diabetes mellitus, recreational drug usage, excessive alcohol usage, and recent adherence to low-carbohydrate diets. All scans were conducted during morning hours, and participants were instructed to arrive after overnight fasting. All individuals were tested for HbA1c, but one participant did not complete the fasting blood glucose measurement as it was part of an oral glucose tolerance test, and they were averse to repeated blood draws over the two hour time course. In the 3T dataset, cognitive status was assessed clinically as cognitively normal, mild cognitive impairment, or dementia. In the cognitively impaired group presented in these analyses, 141 individuals (88 male) had mild cognitive impairment, and 16 individuals (13 male) had dementia. As so few individuals had dementia, the latter two groups were combined into a single group of cognitive impairment in these analyses. In the 7T dataset, all participants were confirmed to be cognitively normal using the CNS Vital Signs neurocognitive test battery (*1*).

#### fMRI analyses

There are a few additional notes regarding specific functional network to aid in interpreting the supplemental figures. Three networks in particular have noticeably fewer (<10 total) ROIs compared to the other functional networks; these are the reward, parieto-medial, and medial temporal networks. Because of this, they are subject to more individual variability in the segregation metric and exhibit noisier, and therefore weaker, associations. Also, the somatomotor dorsal ROIs frequently have overlap with non-brain areas particularly in older brains with volume loss, so the signals from these ROIs are often noisier and must be excluded in the processing pipeline used here, leading to far fewer usable participants in this network. Readers are therefore urged to interpret the results associated with these networks with care, and for this reason these networks largely go un-emphasized in the main text.

#### Selection of control genes for cell density

A potential concern in any spatial transcriptomics approach is that any spatial distribution of pathology is driven simply by cell density; that is, any process could be more correlated with regions where more cells are present. To ensure both targeted and unsupervised analyses presented in this work were not biased by relative cell density, five control genes were selected to provide a range of values for overall cellular density: three markers of neuron density (*RBFOX3, TH, MAP2*) and two markers of glial density (*GFAP, S100B*). *RBFOX3* was selected as a marker of neuron density, as it encodes for NeuN which is typically found in all neurons (*2*). *TH* was selected as an additional marker of neuron density typically associated with interneurons in humans (*3*). *MAP2* was selected as a tertiary marker of neuron density, with a particular emphasis on dendritic density (*5, 6*). *GFAP* was selected as a general marker of (mature) astrocyte density (*6, 7*). *S100B* was selected as a secondary marker of astrocyte density (*8*) and as a marker of oligodendrocyte density (*9*). The relative expression levels of these proteins are shown in the bottom of Fig. 4A to provide a comparison for the main results. They are also listed in the complete list of genes for that figure provided in Table S7, and their exact expression ratios and associated significance levels are provided in Table S9.

#### Gene expression analyses

Regional microarray expression data were obtained from six post-mortem brains (one female, 24– 57 years old, 42.5±3.4) provided by the Allen Human Brain Atlas (*10*) (AHBA, https://human.brain-map.org). Data were processed with the *abagen* toolbox (version 0.1.3; https://github.com/rmarkello/abagen) using a 300-region functionally-defined volumetric atlas in MNI space (*11*). Microarray probes were reannotated using data provided by (*12*); probes not matched to a valid Entrez ID were discarded. Next, probes were filtered based on their expression intensity relative to background noise (*13*), such that probes with intensity less than the background in >=20% of samples across donors were discarded. When multiple probes indexed the expression of the same gene, the probe with the most consistent pattern of regional variation across donors (i.e., differential stability; (*14*)) was selected, calculated with:

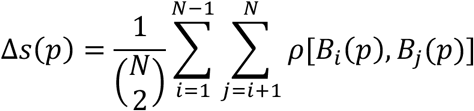

Where *ρ* is Spearman’s rank correlation of the expression of a single probe *p* across regions in two donor brains *B_i_* and *B_j_* and *N*) is the total number of donors. Regions correspond to the structural designations provided in the ontology from the AHBA.

The MNI coordinates of tissue samples were updated to those generated via non-linear registration using the Advanced Normalization Tools (ANTs (*15*); https://github.com/chrisfilo/alleninf). Samples were assigned to brain regions in the provided atlas if their MNI coordinates were within 2mm of a given parcel. To reduce the potential for misassignment, sample-to-region matching was constrained by hemisphere and gross structural divisions (i.e., cortex, subcortex/brainstem, and cerebellum; (*12*)). All tissue samples not assigned to a brain region were discarded. Additionally, due to the inherent limitations of the AHBA in normalizing across subcortical and cortical samples (*16*), ROIs that fell in deep or subcortical or cerebellar regions were omitted from further analyses (see Figure S1 for all ROIs). This selection step was done by using only functional ROIs that overlapped with cortical structures in the Automated Anatomical Labeling atlas (*17*).

Inter-subject variation was addressed by normalizing tissue sample expression values across genes using a robust sigmoid function (*18*):

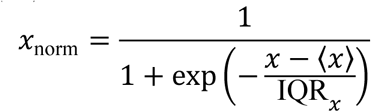

where 〈*x*〉 is the median and IQR*_x_* is the normalized interquartile range of the expression of a single tissue sample across genes. Normalized expression values were then rescaled to the unit interval:

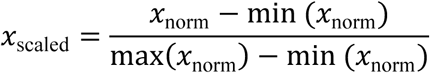

Gene expression values were then normalized across tissue samples using an identical procedure. Samples assigned to the same brain region were averaged separately for each donor and then across donors.

#### Statistical analyses

There is one cosmetic note for statistics in the supplemental sections: in Tables S1-S7, p = 0 means that p < 1e-4. This choice was made to reduce the size of the tables to be more legible as very small p-values are uninformative and are typically tied to the size of the dataset. All p-values greater than this cutoff are explicitly written in the tables.

## Supplementary Figures

**Figure S1:**
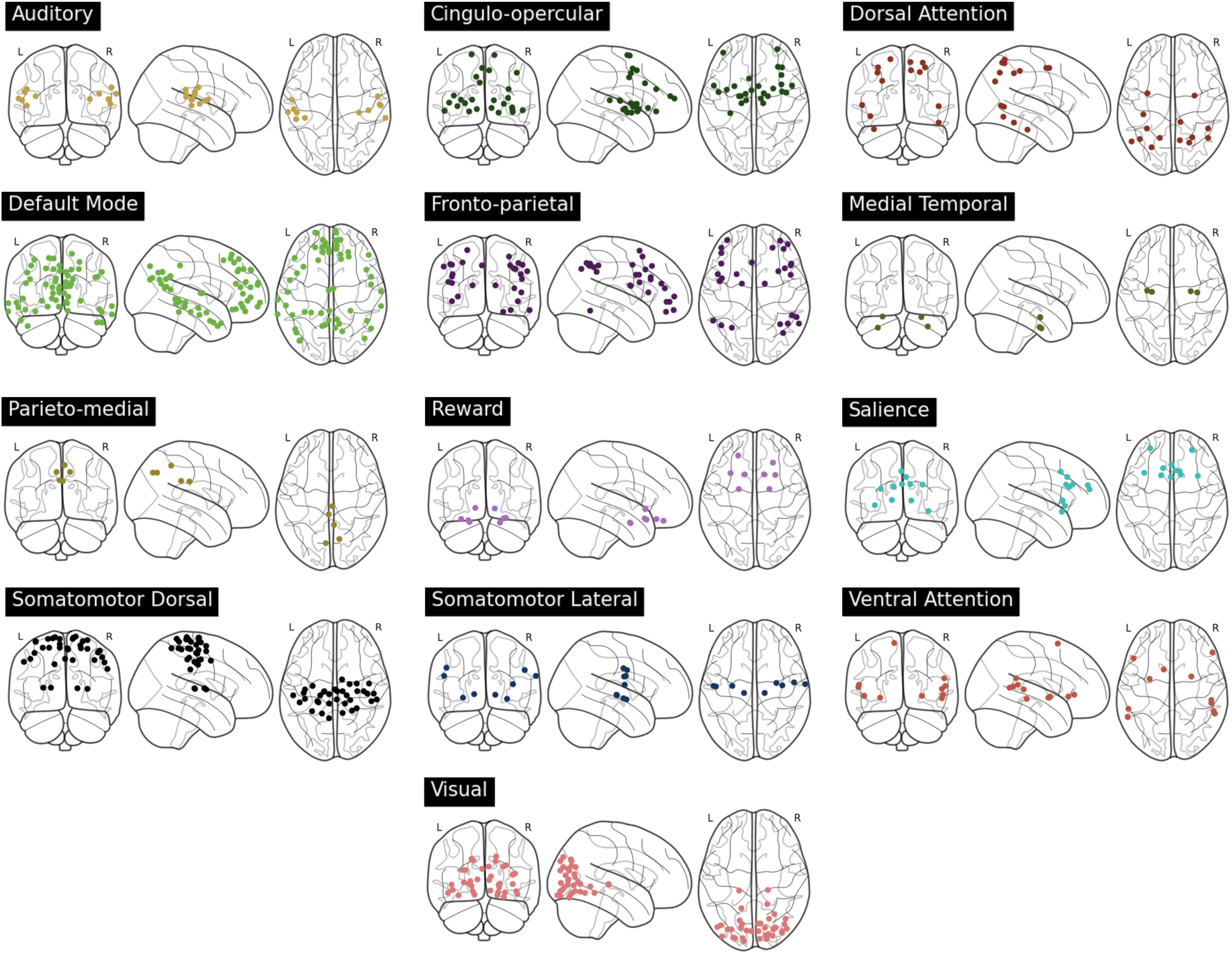
Spatial visualizations for all networks within the atlas used for the analyses presented here. Certain ROIs present in the original atlas have been excluded from these analyses; these include ROIs not assigned to a specific network (12 ROIs total) and ROIs that lie in the cerebellum (26 ROIs total).

**Figure S2:**
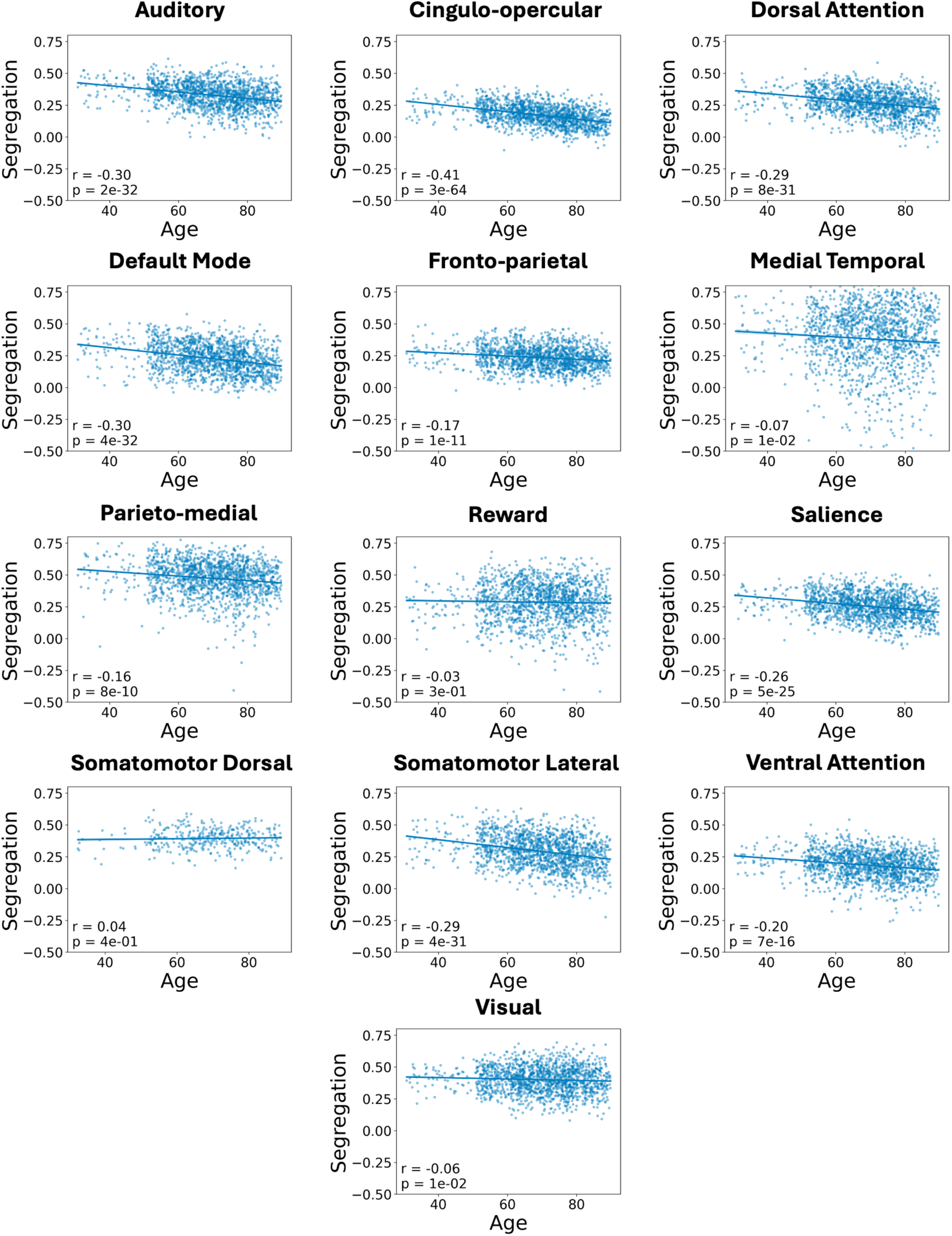
Loss of functional network segregation is associated with age throughout the brain when examined with 3T fMRI. This figure extends Fig. 1A to all the networks examined in these analyses.

**Figure S3:**
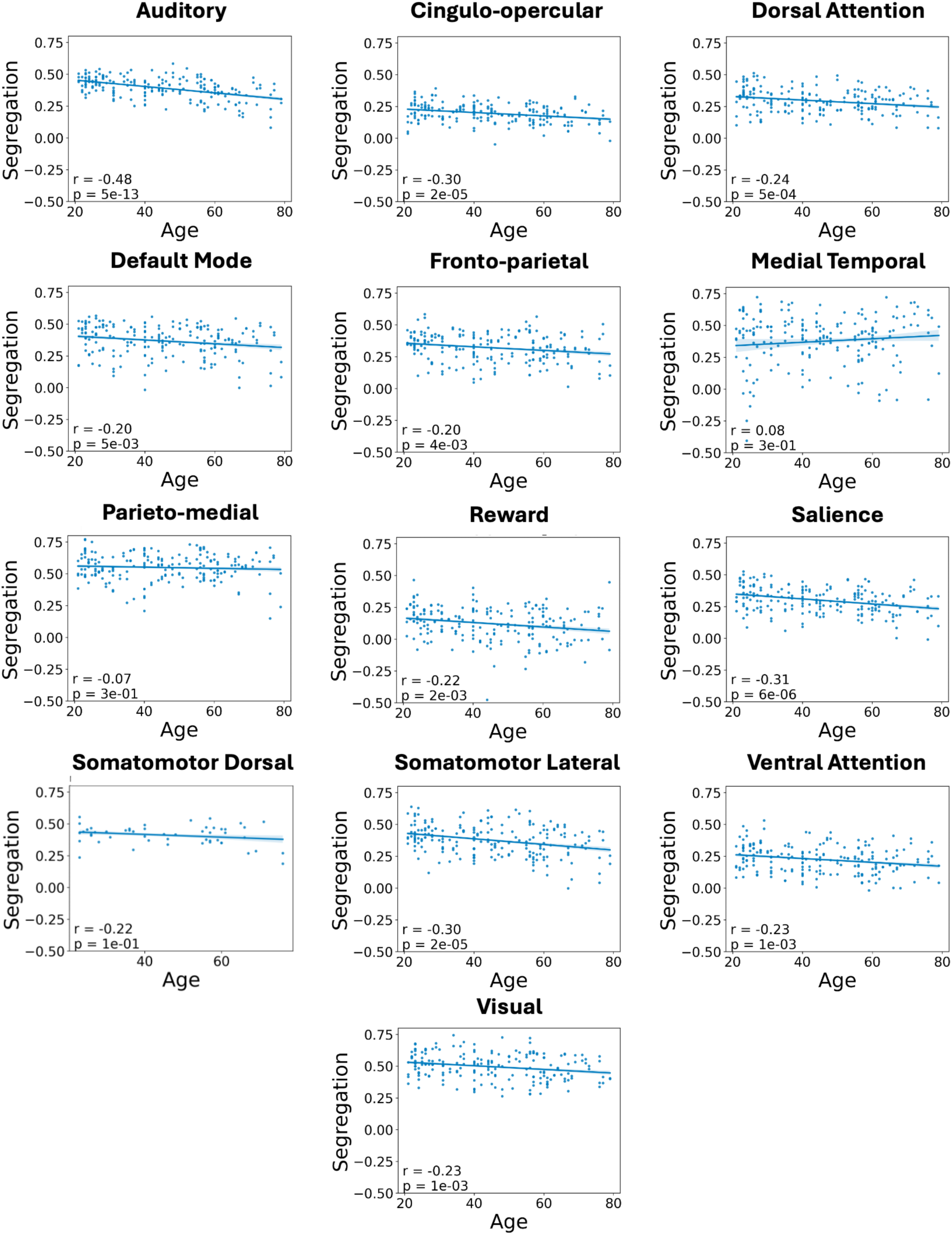
Loss of functional network segregation is associated with age throughout the brain when examined with 7T fMRI. This figure extends Fig. 1B to all the networks examined in these analyses.

**Figure S4:**
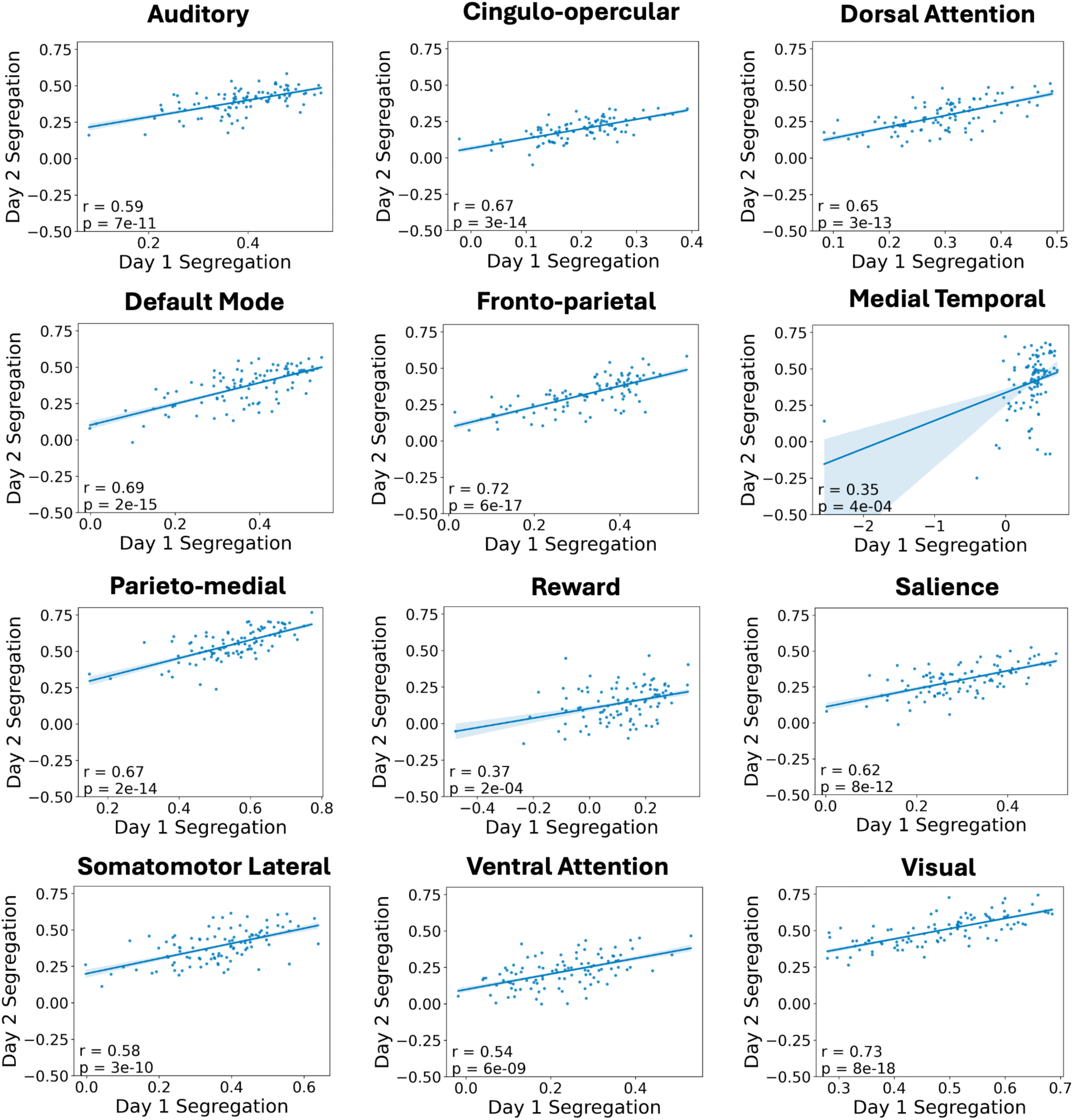
Network segregation is highly correlated within individuals across different scanning days. The relatively high degree of correlation The somatomotor dorsal network is excluded from these results as after removal of low-quality scans for those ROIs there are too few datapoints for a reliable indicator of robustness (see supplemental fMRI details for further discussion).

**Figure S5:**
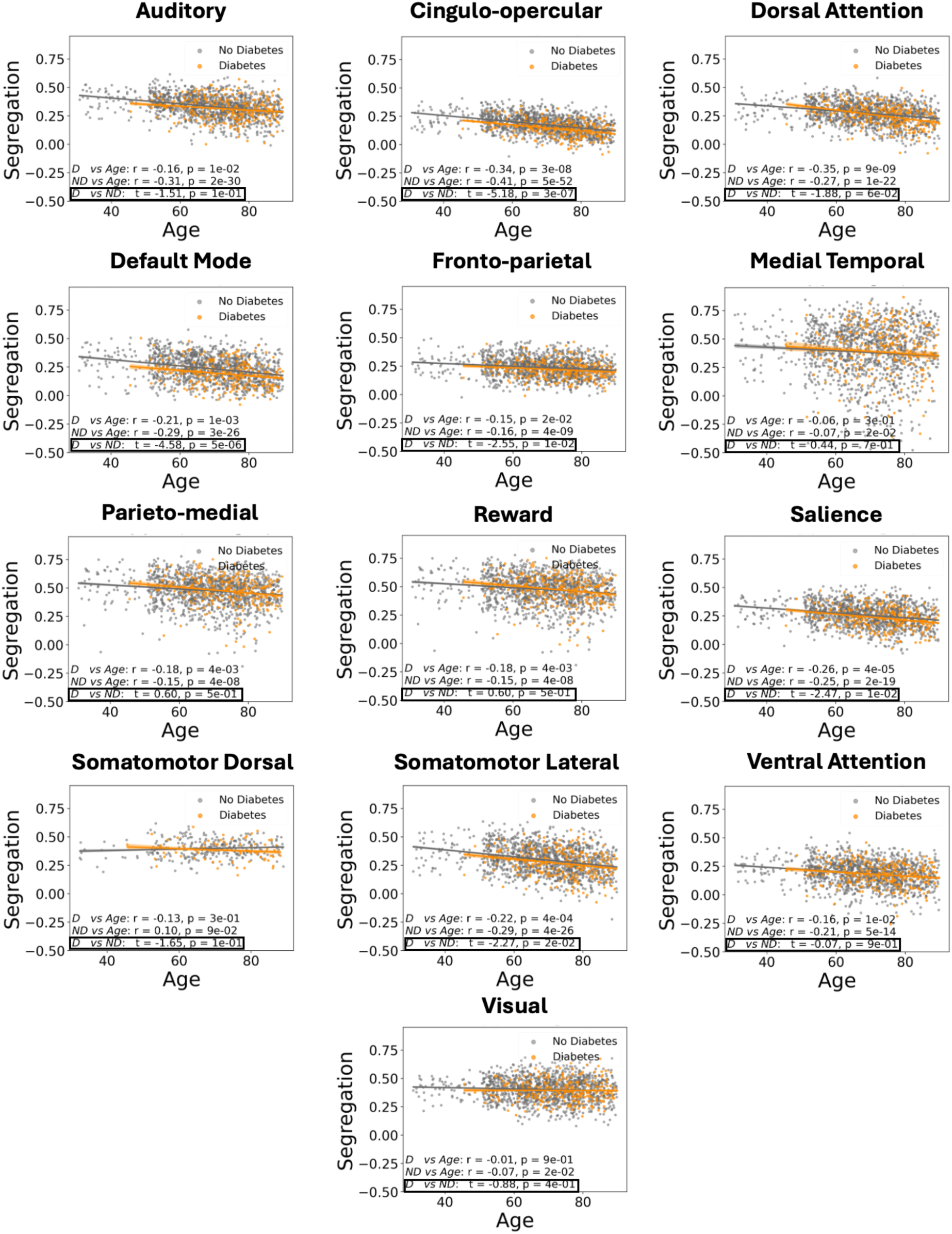
Diabetes is associated with spatially distinct network segregation loss beyond aging. Networks especially affected by diabetes are cingulo-opercular, default mode, fronto-parietal, salience, and somatomotor lateral networks.

**Figure S6:**
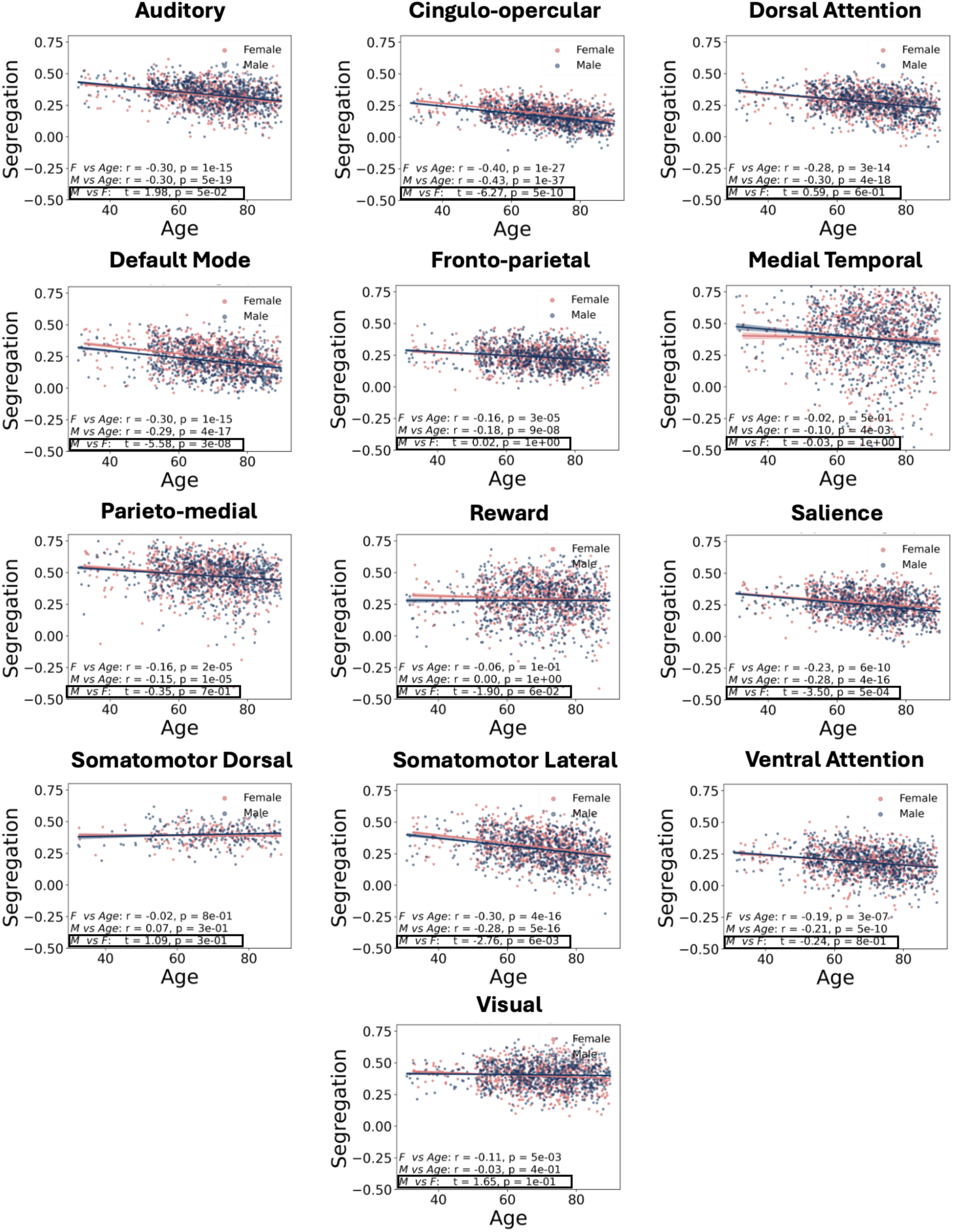
Male sex is associated with similarly spatially distinct network segregation loss beyond aging in the 3T dataset. Cingulo-opercular, default mode, salience, and somatomotor lateral networks were associated with lowered network segregation in male participants in this dataset. There is a marginally increased network segregation in males in the auditory network.

**Figure S7:**
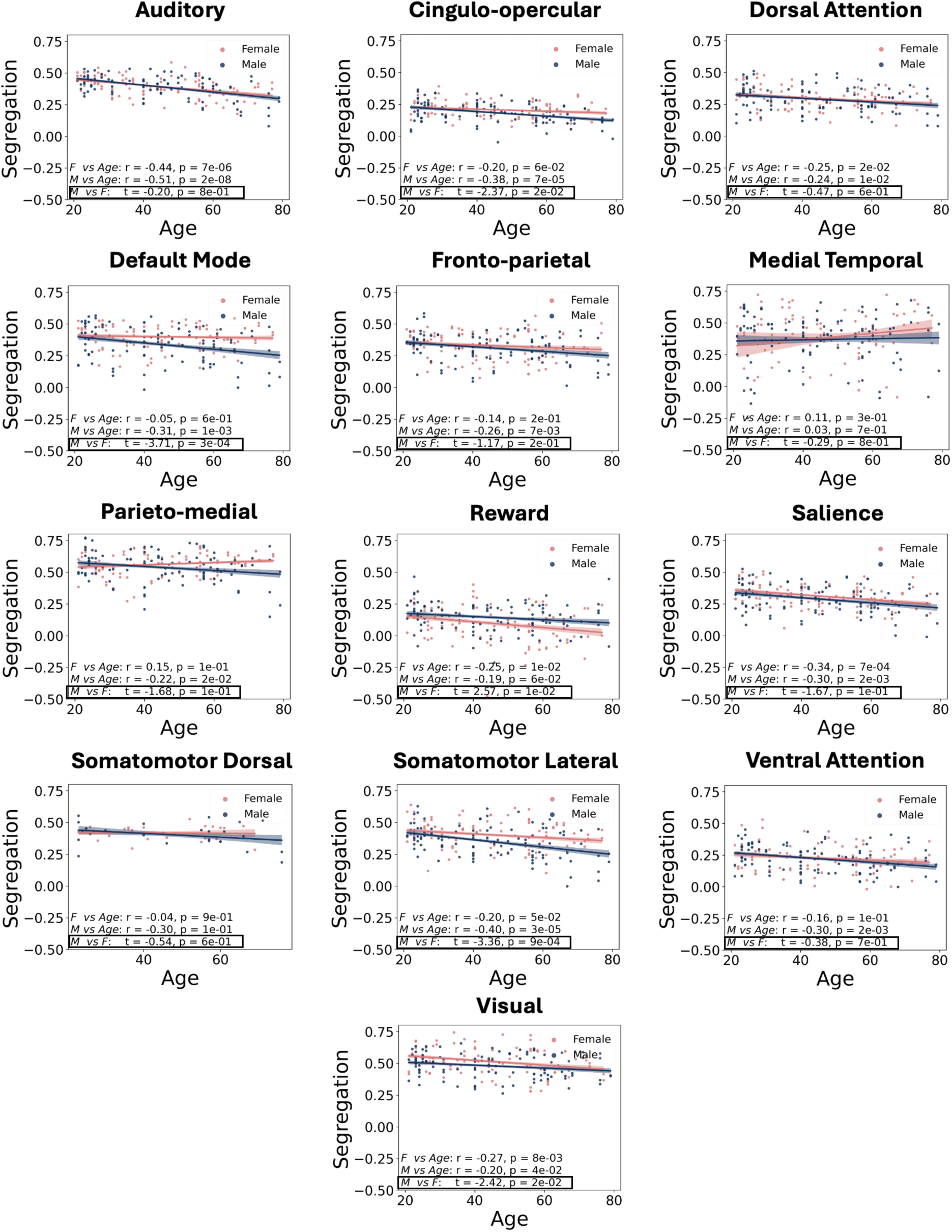
Male sex is associated with similarly spatially distinct network segregation loss beyond aging in the 7T dataset. Cingulo-opercular, default mode, salience, and somatomotor lateral networks were associated with lowered network segregation in male participants in this dataset.

**Figure S8:**
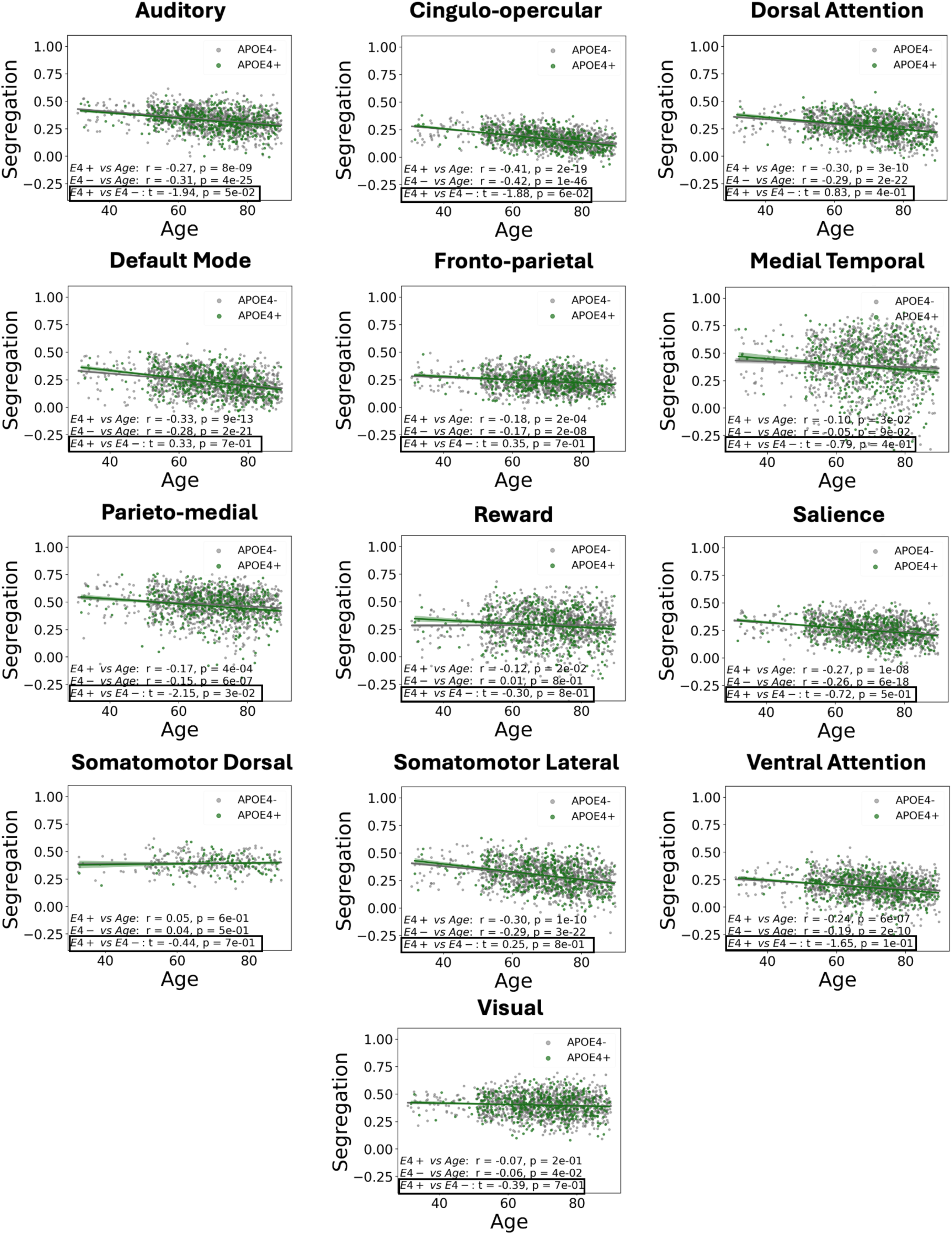
APOE4+ status is associated with network segregation loss beyond aging in different networks than those affected by diabetes and male sex. APOE4+ status is associated with loss of functional segregation in the parieto-medial network, and marginally in the auditory network as well.

## Supplementary Tables

These are the captions associated with each of the supplemental tables. As they are all substantially larger than a standard page, they are included as separate Excel files for ease of viewing. The associated file names are given at the end of each captions.

**Table S1:** Associations of age effects with network segregation in the 7T dataset, corrected for repeated scans of the same individuals. File: TableS1.xlsx.

**Table S2:** Associations of diabetes effects with network segregation in the 3T dataset, with age included in the same OLS model. File: TableS2.xlsx.

**Table S3:** Associations of sex effects with network segregation in both 3T and 7T datasets, with age included in the same OLS models. File: TableS3.xlsx.

**Table S4:** Associations of diabetes and sex with network segregation in the 3T dataset, probing both individual effects and a diabetes*sex interaction effect, with age included in the same OLS model. File: TableS4.xlsx.

**Table S5:** Associations of fasting blood glucose, HbA1c and sex with network segregation in both the 3T and 7T datasets, probing both individual effects and combined glucose*sex or HbA1c*sex interaction effects, with age included in the same OLS models. File: TableS5.xlsx.

**Table S6:** Associations of APOE4 effects with network segregation in the 3T dataset, with age included in the same OLS model. File: TableS6.xlsx.

**Table S7:** Complete list of genes selected for the comparisons shown in Fig. 4A and their respective functional families. File: TableS7.xlsx.

**Table S8:** Entire GSEA results for 500 genes most significantly overexpressed in the diabetes and sex-related functional networks, showing the 10 gene sets that had the most genes overlapping with the top 500. File: TableS8.tsv.

**Table S9:** Full list of all relative expression ratios and p-values (both FDR-corrected as shown in Fig. 4B-C and uncorrected for reference) for all 14,609 genes analyzed. File: TableS9.xlsx.

## Notes

### Competing Interest Statement

The authors have declared no competing interest.

## References

1. N. Bobba-Alves, R.-P. Juster, M. Picard, The energetic cost of allostasis and allostatic load. Psychoneuroendocrinology 146, 105951 (2022).

2. N. Bobba-Alves, G. Sturm, J. Lin, S. A. Ware, K. R. Karan, A. S. Monzel, C. Bris, V. Procaccio, G. Lenaers, A. Higgins-Chen, M. Levine, S. Horvath, B. S. Santhanam, B. A. Kaufman, M. Hirano, E. Epel, M. Picard, Cellular allostatic load is linked to increased energy expenditure and accelerated biological aging. Psychoneuroendocrinology 155, 106322 (2023).

3. T. S. Kraft, V. V. Venkataraman, I. J. Wallace, A. N. Crittenden, N. B. Holowka, J. Stieglitz, J. Harris, D. A. Raichlen, B. Wood, M. Gurven, H. Pontzer, The energetics of uniquely human subsistence strategies. Science 374, eabf0130 (2021).

4. G. M. McKhann, D. S. Knopman, H. Chertkow, B. T. Hyman, C. R. Jack, C. H. Kawas, W. E. Klunk, W. J. Koroshetz, J. J. Manly, R. Mayeux, R. C. Mohs, J. C. Morris, M. N. Rossor, P. Scheltens, M. C. Carrillo, B. Thies, S. Weintraub, C. H. Phelps, The diagnosis of dementia due to Alzheimer’s disease: Recommendations from the National Institute on Aging-Alzheimer’s Association workgroups on diagnostic guidelines for Alzheimer’s disease. Alzheimer’s & Dementia 7, 263–269 (2011).

5. J. T. O’Brien, A. Thomas, Vascular dementia. The Lancet 386, 1698–1706 (2015).

6. N. Villain, M. Fouquet, J.-C. Baron, F. Mézenge, B. Landeau, V. de La Sayette, F. Viader, F. Eustache, B. Desgranges, G. Chételat, Sequential relationships between grey matter and white matter atrophy and brain metabolic abnormalities in early Alzheimer’s disease. Brain 133, 3301– 3314 (2010).

7. A. M. Brickman, F. A. Provenzano, J. Muraskin, J. J. Manly, S. Blum, Z. Apa, Y. Stern, T. R. Brown, J. A. Luchsinger, R. Mayeux, Regional White Matter Hyperintensity Volume, Not Hippocampal Atrophy, Predicts Incident Alzheimer Disease in the Community. Archives of Neurology 69, 1621–1627 (2012).

8. J. Lee, B. J. Burkett, H.-K. Min, M. L. Senjem, E. Dicks, N. Corriveau-Lecavalier, C. T. Mester, H. J. Wiste, E. S. Lundt, M. E. Murray, A. T. Nguyen, R. R. Reichard, H. Botha, J. Graff-Radford, L. R. Barnard, J. L. Gunter, C. G. Schwarz, K. Kantarci, D. S. Knopman, B. F. Boeve, V. J. Lowe, R. C. Petersen, C. R. Jack, D. T. Jones, Synthesizing images of tau pathology from cross-modal neuroimaging using deep learning. Brain 147, 980–995 (2023).

9. C. R. Jack, D. S. Knopman, W. J. Jagust, R. C. Petersen, M. W. Weiner, P. S. Aisen, L. M. Shaw, P. Vemuri, H. J. Wiste, S. D. Weigand, T. G. Lesnick, V. S. Pankratz, M. C. Donohue, J. Q. Trojanowski, Update on hypothetical model of Alzheimer’s disease biomarkers. Lancet Neurol 12, 207–216 (2013).

10. B. Antal, L. P. McMahon, S. F. Sultan, A. Lithen, D. J. Wexler, B. Dickerson, E.-M. Ratai, L. R. Mujica-Parodi, P. Kapahi, C. Isales, Eds. Type 2 diabetes mellitus accelerates brain aging and cognitive decline: Complementary findings from UK Biobank and meta-analyses. eLife 11, e73138 (2022).

11. C. Weistuch, L. R. Mujica-Parodi, R. M. Razban, B. Antal, H. van Nieuwenhuizen, A. Amgalan, K. A. Dill, Metabolism modulates network synchrony in the aging brain. Proceedings of the National Academy of Sciences 118, e2025727118 (2021).

12. L. R. Mujica-Parodi, A. Amgalan, S. F. Sultan, B. Antal, X. Sun, S. Skiena, A. Lithen, N. Adra, E.-M. Ratai, C. Weistuch, S. T. Govindarajan, H. H. Strey, K. A. Dill, S. M. Stufflebeam, R. L. Veech, K. Clarke, Diet modulates brain network stability, a biomarker for brain aging, in young adults. Proceedings of the National Academy of Sciences 117, 6170–6177 (2020).

13. S. S. Haas, F. Abbasi, K. Watson, T. Robakis, A. Myoraku, S. Frangou, N. Rasgon, Metabolic Status Modulates Global and Local Brain Age Estimates in Overweight and Obese Adults. Biological Psychiatry: Cognitive Neuroscience and Neuroimaging (2024), doi:10.1016/j.bpsc.2024.11.017.

14. D. Tomasi, N. D. Volkow, Aging and Functional Brain Networks. Mol Psychiatry 17, 471– 558 (2012).

15. P. Manza, C. E. Wiers, E. Shokri-Kojori, D. Kroll, D. Feldman, M. Schwandt, G.-J. Wang, D. Tomasi, N. D. Volkow, Brain Network Segregation and Glucose Energy Utilization: Relevance for Age-Related Differences in Cognitive Function. Cereb Cortex 30, 5930–5942 (2020).

16. B. B. Antal, H. van Nieuwenhuizen, A. G. Chesebro, H. H. Strey, D. T. Jones, K. Clarke, C. Weistuch, E.-M. Ratai, K. A. Dill, L. R. Mujica-Parodi, Brain aging shows nonlinear transitions, suggesting a midlife “critical window” for metabolic intervention. Proceedings of the National Academy of Sciences 122, e2416433122 (2025).

17. N. Scarmeas, J. A. Luchsinger, N. Schupf, A. M. Brickman, S. Cosentino, M. X. Tang, Y. Stern, Physical Activity, Diet, and Risk of Alzheimer Disease. JAMA 302, 627–637 (2009).

18. S. M. de la Monte, J. R. Wands, Alzheimer’s Disease Is Type 3 Diabetes–Evidence Reviewed. J Diabetes Sci Technol 2, 1101–1113 (2008).

19. S. Chatterjee, S. A. E. Peters, M. Woodward, S. Mejia Arango, G. D. Batty, N. Beckett, A. Beiser, A. R. Borenstein, P. K. Crane, M. Haan, L. B. Hassing, K. M. Hayden, Y. Kiyohara, E. B. Larson, C.-Y. Li, T. Ninomiya, T. Ohara, R. Peters, T. C. Russ, S. Seshadri, B. H. Strand, R. Walker, W. Xu, R. R. Huxley, Type 2 Diabetes as a Risk Factor for Dementia in Women Compared With Men: A Pooled Analysis of 2.3 Million People Comprising More Than 100,000 Cases of Dementia. Diabetes Care 39, 300–307 (2016).

20. N. Ho, M. S. Sommers, I. Lucki, Effects of diabetes on hippocampal neurogenesis: Links to cognition and depression. Neuroscience & Biobehavioral Reviews 37, 1346–1362 (2013).

21. A. B. Kaiser, N. Zhang, W. V. Der Pluijm, Global Prevalence of Type 2 Diabetes over the Next Ten Years (2018-2028). Diabetes 67, 202-LB (2018).

22. D. A. Levine, A. L. Gross, E. M. Briceño, N. Tilton, B. J. Giordani, J. B. Sussman, R. A. Hayward, J. F. Burke, S. Hingtgen, M. S. V. Elkind, J. J. Manly, R. F. Gottesman, D. J. Gaskin, S. Sidney, R. L. Sacco, S. E. Tom, C. B. Wright, K. Yaffe, A. T. Galecki, Sex Differences in Cognitive Decline Among US Adults. JAMA Network Open 4, e210169 (2021).

23. C. Barth, A. Crestol, A.-M. G. de Lange, L. A. M. Galea, Sex steroids and the female brain across the lifespan: insights into risk of depression and Alzheimer’s disease. The Lancet Diabetes & Endocrinology 11, 926–941 (2023).

24. J. K. Russell, C. K. Jones, P. A. Newhouse, The Role of Estrogen in Brain and Cognitive Aging. Neurotherapeutics 16, 649–665 (2019).

25. A. R. Genazzani, N. Pluchino, S. Luisi, M. Luisi, Estrogen, cognition and female ageing. Human Reproduction Update 13, 175–187 (2007).

26. R. N. M. Saleh, M. Hornberger, C. W. Ritchie, A. M. Minihane, Hormone replacement therapy is associated with improved cognition and larger brain volumes in at-risk APOE4 women: results from the European Prevention of Alzheimer’s Disease (EPAD) cohort. Alzheimer’s Research & Therapy 15, 10 (2023).

27. S. Oveisgharan, Z. Arvanitakis, L. Yu, J. Farfel, J. A. Schneider, D. A. Bennett, Sex differences in Alzheimer’s disease and common neuropathologies of aging. Acta Neuropathol 136, 887–900 (2018).

28. A. Torres-Espin, H. L. Radabaugh, S. Treiman, S. S. Fitzsimons, D. Harvey, A. Chou, C. A. Lindbergh, K. B. Casaletto, L. Goldberger, A. M. Staffaroni, P. Maillard, B. L. Miller, C. DeCarli, J. D. Hinman, A. R. Ferguson, J. H. Kramer, F. M. Elahi, Sexually dimorphic differences in angiogenesis markers are associated with brain aging trajectories in humans. Science Translational Medicine 16, eadk3118 (2024).

29. C. Lopez-Lee, E. R. S. Torres, G. Carling, L. Gan, Mechanisms of sex differences in Alzheimer’s disease. Neuron 112, 1208–1221 (2024).

30. M. Y. Chan, D. C. Park, N. K. Savalia, S. E. Petersen, G. S. Wig, Decreased segregation of brain systems across the healthy adult lifespan. Proceedings of the National Academy of Sciences 111, E4997–E5006 (2014).

31. M. J. Hawrylycz, E. S. Lein, A. L. Guillozet-Bongaarts, E. H. Shen, L. Ng, J. A. Miller, L. N. van de Lagemaat, K. A. Smith, A. Ebbert, Z. L. Riley, C. Abajian, C. F. Beckmann, A. Bernard, D. Bertagnolli, A. F. Boe, P. M. Cartagena, M. M. Chakravarty, M. Chapin, J. Chong, R. A. Dalley, B. D. Daly, C. Dang, S. Datta, N. Dee, T. A. Dolbeare, V. Faber, D. Feng, D. R. Fowler, J. Goldy, B. W. Gregor, Z. Haradon, D. R. Haynor, J. G. Hohmann, S. Horvath, R. E. Howard, A. Jeromin, J. M. Jochim, M. Kinnunen, C. Lau, E. T. Lazarz, C. Lee, T. A. Lemon, L. Li, Y. Li, J. A. Morris, C. C. Overly, P. D. Parker, S. E. Parry, M. Reding, J. J. Royall, J. Schulkin, P. A. Sequeira, C. R. Slaughterbeck, S. C. Smith, A. J. Sodt, S. M. Sunkin, B. E. Swanson, M. P. Vawter, D. Williams, P. Wohnoutka, H. R. Zielke, D. H. Geschwind, P. R. Hof, S. M. Smith, C. Koch, S. G. N. Grant, A. R. Jones, An anatomically comprehensive atlas of the adult human brain transcriptome. Nature 489, 391–399 (2012).

32. B. A. Seitzman, C. Gratton, S. Marek, R. V. Raut, N. U. F. Dosenbach, B. L. Schlaggar, S. E. Petersen, D. J. Greene, A set of functionally-defined brain regions with improved representation of the subcortex and cerebellum. NeuroImage 206, 116290 (2020).

33. A. Altmann, L. Tian, V. W. Henderson, M. D. Greicius, Sex Modifies the APOE-Related Risk of Developing Alzheimer’s Disease. Ann Neurol 75, 563–573 (2014).

34. S. Walters, A. G. Contreras, J. M. Eissman, S. Mukherjee, M. L. Lee, S.-E. Choi, P. Scollard, E. H. Trittschuh, J. B. Mez, W. S. Bush, B. W. Kunkle, A. C. Naj, A. Peterson, K. A. Gifford, M. L. Cuccaro, C. Cruchaga, M. A. Pericak-Vance, L. A. Farrer, L.-S. Wang, J. L. Haines, A. L. Jefferson, W. A. Kukull, C. D. Keene, A. J. Saykin, P. M. Thompson, E. R. Martin, D. A. Bennett, L. L. Barnes, J. A. Schneider, P. K. Crane, T. J. Hohman, L. Dumitrescu, A. D. G. C. Alzheimer’s Disease Neuroimaging Initiative and Alzheimer’s Disease Sequencing Project, Associations of Sex, Race, and Apolipoprotein E Alleles With Multiple Domains of Cognition Among Older Adults. JAMA Neurology 80, 929–939 (2023).

35. M. E. Wood, L. Y. Xiong, Y. Y. Wong, R. F. Buckley, W. Swardfager, M. Masellis, A. S. P. Lim, E. Nichols, R. La Joie, K. B. Casaletto, R. G. Kumar, K. Dams-O’Connor, P. Palta, K. M. George, C. L. Satizabal, L. L. Barnes, J. A. Schneider, A. P. Binet, S. Villeneuve, J. Pa, A. M. Brickman, S. E. Black, J. S. Rabin, Sex differences in associations between APOE ε2 and longitudinal cognitive decline. Alzheimers Dement 19, 4651–4661 (2023).

36. S. Yan, C. Zheng, M. D. Paranjpe, Y. Li, W. Li, X. Wang, T. L. S. Benzinger, J. Lu, Y. Zhou, for the Alzheimer’s Disease Neuroimaging Initiative, Sex modifies APOE ε4 dose effect on brain tau deposition in cognitively impaired individuals. Brain 144, 3201–3211 (2021).

37. G. Ashrafi, Z. Wu, R. J. Farrell, T. A. Ryan, GLUT4 Mobilization Supports Energetic Demands of Active Synapses. Neuron 93, 606–615.e3 (2017).

38. B. Kula, B. Antal, C. Weistuch, F. Gackière, A. Barre, V. Velado, J. M. Hubbard, M. Kukley, L. R. Mujica-Parodi, N. A. Smith, D-ꞵ-hydroxybutyrate stabilizes hippocampal CA3-CA1 circuit during acute insulin resistance. PNAS Nexus 3, pgae196 (2024).

39. M. Namwanje, C. W. Brown, Activins and Inhibins: Roles in Development, Physiology, and Disease. Cold Spring Harb Perspect Biol 8, a021881 (2016).

40. P. Dewing, M. I. Boulware, K. Sinchak, A. Christensen, P. G. Mermelstein, P. Micevych, Membrane Estrogen Receptor-α Interactions with Metabotropic Glutamate Receptor 1a Modulate Female Sexual Receptivity in Rats. J. Neurosci. 27, 9294–9300 (2007).

41. D. Grove-Strawser, M. I. Boulware, P. G. Mermelstein, Membrane Estrogen Receptors Activate the Metabotropic Glutamate Receptors mGluR5 and mGluR3 to Bidirectionally Regulate CREB Phosphorylation in Female Rat Striatal Neurons. Neuroscience 170, 1045–1055 (2010).

42. M. P. Fagan, D. Ameroso, A. Meng, A. Rock, J. Maguire, M. Rios, Essential and sex-specific effects of mGluR5 in ventromedial hypothalamus regulating estrogen signaling and glucose balance. Proceedings of the National Academy of Sciences 117, 19566–19577 (2020).

43. A. Subramanian, P. Tamayo, V. K. Mootha, S. Mukherjee, B. L. Ebert, M. A. Gillette, A. Paulovich, S. L. Pomeroy, T. R. Golub, E. S. Lander, J. P. Mesirov, Gene set enrichment analysis: A knowledge-based approach for interpreting genome-wide expression profiles. Proceedings of the National Academy of Sciences 102, 15545–15550 (2005).

44. R. T. Pillich, J. Chen, C. Churas, S. Liu, K. Ono, D. Otasek, D. Pratt, NDEx: Accessing Network Models and Streamlining Network Biology Workflows. Current Protocols 1, e258 (2021).

45. M. Filippi, C. Cividini, S. Basaia, E. G. Spinelli, V. Castelnovo, M. Leocadi, E. Canu, F. Agosta, Age-related vulnerability of the human brain connectome. Mol Psychiatry 28, 5350– 5358 (2023).

46. G. Argiris, Y. Stern, C. Habeck, Cross-sectional and Longitudinal Age-related Disintegration in Functional Connectivity: Reference Ability Neural Network Cohort. J Cogn Neurosci 36, 2045–2066 (2024).

47. H. K. Ballard, T. B. Jackson, A. C. Symm, T. H. Hicks, J. A. Bernard, Age-related differences in functional network segregation in the context of sex and reproductive stage. Hum Brain Mapp 44, 1949–1963 (2022).

48. W. Stanford, P. J. Mucha, E. Dayan, Age-related differences in network controllability are mitigated by redundancy in large-scale brain networks. Commun Biol 7, 701 (2024).

49. R. M. Razban, B. B. Antal, K. A. Dill, L. R. Mujica-Parodi, Brain signaling becomes less integrated and more segregated with age. Network Neuroscience, 1–36 (2024).

50. H. A. Deery, E. X. Liang, M. N. Siddiqui, G. Murray, K. Voigt, R. Di Paolo, C. Moran, G. F. Egan, S. D. Jamadar, Reconfiguration of metabolic connectivity in ageing. Commun Biol 7, 1–14 (2024).

51. T. Stoica, L. K. Knight, F. Naaz, S. C. Patton, B. E. Depue, Gender differences in functional connectivity during emotion regulation. Neuropsychologia 156, 107829 (2021).

52. B. A. Zielinski, D. S. Andrews, J. K. Lee, M. Solomon, S. J. Rogers, B. Heath, C. W. Nordahl, D. G. Amaral, Sex-dependent structure of socioemotional salience, executive control, and default mode networks in preschool-aged children with autism. NeuroImage 257, 119252 (2022).

53. D. T. Jones, D. S. Knopman, J. L. Gunter, J. Graff-Radford, P. Vemuri, B. F. Boeve, R. C. Petersen, M. W. Weiner, C. R. Jack Jr, on behalf of the Alzheimer’s Disease Neuroimaging Initiative, Cascading network failure across the Alzheimer’s disease spectrum. Brain 139, 547– 562 (2016).

54. D. T. Jones, M. M. Machulda, P. Vemuri, E. M. McDade, G. Zeng, M. L. Senjem, J. L. Gunter, S. A. Przybelski, R. T. Avula, D. S. Knopman, B. F. Boeve, R. C. Petersen, C. R. Jack, Age-related changes in the default mode network are more advanced in Alzheimer disease. Neurology 77, 1524–1531 (2011).

55. M. Habes, A. M. Jacobson, B. H. Braffett, T. Rashid, C. M. Ryan, H. Shou, Y. Cui, C. Davatzikos, J. A. Luchsinger, G. J. Biessels, I. Bebu, R. A. Gubitosi-Klug, R. N. Bryan, I. M. Nasrallah, DCCT/EDIC Research Group, Patterns of Regional Brain Atrophy and Brain Aging in Middle- and Older-Aged Adults With Type 1 Diabetes. JAMA Network Open 6, e2316182 (2023).

56. S. E. Choi, B. Roy, M. Freeby, R. Mullur, M. A. Woo, R. Kumar, Prefrontal Cortex Brain Damage and Glycemic Control in Patients with Type 2 Diabetes. J Diabetes 12, 465–473 (2020).

57. A. Backeström, K. Papadopoulos, S. Eriksson, T. Olsson, M. Andersson, K. Blennow, H. Zetterberg, L. Nyberg, O. Rolandsson, Acute hyperglycaemia leads to altered frontal lobe brain activity and reduced working memory in type 2 diabetes. PLoS One 16, e0247753 (2021).

58. E. E. Sundermann, M. Tran, P. M. Maki, M. W. Bondi, Sex differences in the association between apolipoprotein E ε4 allele and Alzheimer’s disease markers. Alzheimers Dement (Amst) 10, 438–447 (2018).

59. J. S. Damoiseaux, W. W. Seeley, J. Zhou, W. R. Shirer, G. Coppola, A. Karydas, H. J. Rosen, B. L. Miller, J. H. Kramer, M. D. Greicius, Gender Modulates the APOE ε4 Effect in Healthy Older Adults: Convergent Evidence from Functional Brain Connectivity and Spinal Fluid Tau Levels. J. Neurosci. 32, 8254–8262 (2012).

60. I. C. Turney, A. G. Chesebro, M. A. Rentería, P. J. Lao, J. M. Beato, N. Schupf, R. Mayeux, J. J. Manly, A. M. Brickman, APOE ε4 and resting-state functional connectivity in racially/ethnically diverse older adults. Alzheimer’s & Dementia: Diagnosis, Assessment & Disease Monitoring 12, e12094 (2020).

61. M. M. Machulda, D. T. Jones, P. Vemuri, E. McDade, R. Avula, S. Przybelski, B. F. Boeve, D. S. Knopman, R. C. Petersen, C. R. Jack Jr, Effect of APOE ε4 Status on Intrinsic Network Connectivity in Cognitively Normal Elderly Subjects. Archives of Neurology 68, 1131–1136 (2011).

62. P. Geraldes, J. Hiraoka-Yamamoto, M. Matsumoto, A. Clermont, M. Leitges, A. Marette, L. P. Aiello, T. S. Kern, G. L. King, Activation of PKCδ and SHP1 by hyperglycemia causes vascular cell apoptosis and diabetic retinopathy. Nat Med 15, 1298–1306 (2009).

63. G. Lu, J. Li, T. Gao, Q. Liu, O. Chen, X. Zhang, M. Xiao, Y. Guo, J. Wang, Y. Tang, J. Gu, Integration of dietary nutrition and TRIB3 action into diabetes mellitus. Nutrition Reviews 82, 361–373 (2024).

64. A. H. K. Fok, Y. Huang, B. W. L. So, Q. Zheng, C. S. C. Tse, X. Li, K. K.-Y. Wong, J. Huang, K.-O. Lai, C. S. W. Lai, KIF5B plays important roles in dendritic spine plasticity and dendritic localization of PSD95 and FMRP in the mouse cortex *in vivo*. Cell Reports 43, 113906 (2024).

65. J. Zhao, A. H. K. Fok, R. Fan, P.-Y. Kwan, H.-L. Chan, L. H.-Y. Lo, Y.-S. Chan, W.-H. Yung, J. Huang, C. S. W. Lai, K.-O. Lai, E. Kim, J. A. Cooper, L. Bao, Eds. Specific depletion of the motor protein KIF5B leads to deficits in dendritic transport, synaptic plasticity and memory. eLife 9, e53456 (2020).

66. S. Semiz, J. G. Park, S. M. C. Nicoloro, P. Furcinitti, C. Zhang, A. Chawla, J. Leszyk, M. P. Czech, Conventional kinesin KIF5B mediates insulin-stimulated GLUT4 movements on microtubules. The EMBO Journal 22, 2387–2399 (2003).

67. E. Palomer, N. Martín-Flores, S. Jolly, P. Pascual-Vargas, S. Benvegnù, M. Podpolny, S. Teo, K. Vaher, T. Saito, T. C. Saido, P. Whiting, P. C. Salinas, Epigenetic repression of Wnt receptors in AD: a role for Sirtuin2-induced H4K16ac deacetylation of Frizzled1 and Frizzled7 promoters. Molecular Psychiatry 27, 3024 (2022).

68. Z. Pei, Y. He, J. C. Bean, Y. Yang, H. Liu, M. Yu, K. Yu, I. Hyseni, X. Cai, H. Liu, N. Qu, L. Tu, K. M. Conde, M. Wang, Y. Li, N. Yin, N. Zhang, J. Han, C. HS. Potts, N. A. Scarcelli, Z. Yan, P. Xu, Q. Wu, Y. He, Y. Xu, C. Wang, Gabra5 plays a sexually dimorphic role in POMC neuron activity and glucose balance. Front Endocrinol (Lausanne) 13, 889122 (2022).

69. A. R. Rau, S. T. Hentges, The Relevance of AgRP Neuron-Derived GABA Inputs to POMC Neurons Differs for Spontaneous and Evoked Release. J Neurosci 37, 7362–7372 (2017).

70. D. Trabzuni, A. Ramasamy, S. Imran, R. Walker, C. Smith, M. E. Weale, J. Hardy, M. Ryten, Widespread sex differences in gene expression and splicing in the adult human brain. Nat Commun 4, 2771 (2013).

71. L. Han, M. Y. Chan, P. F. Agres, E. Winter-Nelson, Z. Zhang, G. S. Wig, Measures of resting-state brain network segregation and integration vary in relation to data quantity: implications for within and between subject comparisons of functional brain network organization. Cereb Cortex 34, bhad506 (2024).

72. B. Lehallier, D. Gate, N. Schaum, T. Nanasi, S. E. Lee, H. Yousef, P. M. Losada, D. Berdnik, A. Keller, J. Verghese, S. Sathyan, C. Franceschi, S. Milman, N. Barzilai, T. Wyss-Coray, Undulating changes in human plasma proteome profiles across the lifespan. Nat Med 25, 1843–1850 (2019).

73. X. Shen, C. Wang, X. Zhou, W. Zhou, D. Hornburg, S. Wu, M. P. Snyder, Nonlinear dynamics of multi-omics profiles during human aging. Nat Aging 4, 1619–1634 (2024).

74. R. O. Roberts, Y. E. Geda, D. S. Knopman, R. H. Cha, V. S. Pankratz, B. F. Boeve, R. J. Ivnik, E. G. Tangalos, R. C. Petersen, W. A. Rocca, The Mayo Clinic Study of Aging: Design and Sampling, Participation, Baseline Measures and Sample Characteristics. Neuroepidemiology 30, 58–69 (2008).

75. R. Kumar, L. Tan, A. Kriegstein, A. Lithen, J. R. Polimeni, L. R. Mujica-Parodi, H. H. Strey, Ground-truth “resting-state” signal provides data-driven estimation and correction for scanner distortion of fMRI time-series dynamics. Neuroimage 227, 117584 (2021).

76. O. Esteban, C. J. Markiewicz, R. W. Blair, C. A. Moodie, A. I. Isik, A. Erramuzpe, J. D. Kent, M. Goncalves, E. DuPre, M. Snyder, H. Oya, S. S. Ghosh, J. Wright, J. Durnez, R. A. Poldrack, K. J. Gorgolewski, fMRIPrep: a robust preprocessing pipeline for functional MRI. Nat Methods 16, 111–116 (2019).

77. A. Abraham, F. Pedregosa, M. Eickenberg, P. Gervais, A. Mueller, J. Kossaifi, A. Gramfort, B. Thirion, G. Varoquaux, Machine learning for neuroimaging with scikit-learn. Front. Neuroinform. 8 (2014), doi:10.3389/fninf.2014.00014.

78. S. Wang, D. J. Peterson, J. C. Gatenby, W. Li, T. J. Grabowski, T. M. Madhyastha, Evaluation of Field Map and Nonlinear Registration Methods for Correction of Susceptibility Artifacts in Diffusion MRI. Front Neuroinform 11, 17 (2017).

79. M. H. Hagenauer, Y. Sannah, E. K. Hebda-Bauer, C. Rhoads, A. M. O’Connor, E. Flandreau, S. J. Watson, H. Akil, Resource: A curated database of brain-related functional gene sets (Brain.GMT). MethodsX 13, 102788 (2024).

80. R. D. Markello, J. Y. Hansen, Z.-Q. Liu, V. Bazinet, G. Shafiei, L. E. Suárez, N. Blostein, J. Seidlitz, S. Baillet, T. D. Satterthwaite, M. M. Chakravarty, A. Raznahan, B. Misic, neuromaps: structural and functional interpretation of brain maps. Nat Methods 19, 1472–1479 (2022).

81. J. B. Burt, M. Helmer, M. Shinn, A. Anticevic, J. D. Murray, Generative modeling of brain maps with spatial autocorrelation. NeuroImage 220, 117038 (2020).

82. Y. Benjamini, Y. Hochberg, Controlling the False Discovery Rate: A Practical and Powerful Approach to Multiple Testing. Journal of the Royal Statistical Society. Series B (Methodological) 57, 289–300 (1995).

83. A. Liberzon, C. Birger, H. Thorvaldsdóttir, M. Ghandi, J. P. Mesirov, P. Tamayo, The Molecular Signatures Database (MSigDB) hallmark gene set collection. Cell Syst 1, 417–425 (2015).

84. G. Gryglewski, R. Seiger, G. M. James, G. M. Godbersen, A. Komorowski, J. Unterholzner, P. Michenthaler, A. Hahn, W. Wadsak, M. Mitterhauser, S. Kasper, R. Lanzenberger, Spatial analysis and high resolution mapping of the human whole-brain transcriptome for integrative analysis in neuroimaging. Neuroimage 176, 259–267 (2018).

## Supplementary References

1. C. T. Gualtieri, L. G. Johnson, Reliability and validity of a computerized neurocognitive test battery, CNS Vital Signs. Archives of Clinical Neuropsychology 21, 623–643 (2006).

2. V. V. Gusel’nikova, D. E. Korzhevskiy, NeuN As a Neuronal Nuclear Antigen and Neuron Differentiation Marker. Acta Naturae 7, 42–47 (2015).

3. R. Benavides-Piccione, J. DeFelipe, Distribution of neurons expressing tyrosine hydroxylase in the human cerebral cortex. J Anat 211, 212–222 (2007).

4. M. H. Soltani, R. Pichardo, Z. Song, N. Sangha, F. Camacho, K. Satyamoorthy, O. P. Sangueza, V. Setaluri, Microtubule-Associated Protein 2, a Marker of Neuronal Differentiation, Induces Mitotic Defects, Inhibits Growth of Melanoma Cells, and Predicts Metastatic Potential of Cutaneous Melanoma. Am J Pathol 166, 1841–1850 (2005).

5. R. A. DeGiosio, M. J. Grubisha, M. L. MacDonald, B. C. McKinney, C. J. Camacho, R. A. Sweet, More than a marker: potential pathogenic functions of MAP2. Front. Mol. Neurosci. 15 (2022), doi:10.3389/fnmol.2022.974890.

6. J. Middeldorp, E. M. Hol, GFAP in health and disease. Prog Neurobiol 93, 421–443 (2011).

7. M. V. Sofroniew, H. V. Vinters, Astrocytes: biology and pathology. Acta Neuropathol 119, 7– 35 (2010).

8. F. Brozzi, C. Arcuri, I. Giambanco, R. Donato, S100B Protein Regulates Astrocyte Shape and Migration via Interaction with Src Kinase. J Biol Chem 284, 8797–8811 (2009).

9. J. C. Deloulme, E. Raponi, B. J. Gentil, N. Bertacchi, A. Marks, G. Labourdette, J. Baudier, Nuclear expression of S100B in oligodendrocyte progenitor cells correlates with differentiation toward the oligodendroglial lineage and modulates oligodendrocytes maturation. Mol Cell Neurosci 27, 453–465 (2004).

10. M. J. Hawrylycz, E. S. Lein, A. L. Guillozet-Bongaarts, E. H. Shen, L. Ng, J. A. Miller, L. N. van de Lagemaat, K. A. Smith, A. Ebbert, Z. L. Riley, C. Abajian, C. F. Beckmann, A. Bernard, D. Bertagnolli, A. F. Boe, P. M. Cartagena, M. M. Chakravarty, M. Chapin, J. Chong, R. A. Dalley, B. D. Daly, C. Dang, S. Datta, N. Dee, T. A. Dolbeare, V. Faber, D. Feng, D. R. Fowler, J. Goldy, B. W. Gregor, Z. Haradon, D. R. Haynor, J. G. Hohmann, S. Horvath, R. E. Howard, A. Jeromin, J. M. Jochim, M. Kinnunen, C. Lau, E. T. Lazarz, C. Lee, T. A. Lemon, L. Li, Y. Li, J. A. Morris, C. C. Overly, P. D. Parker, S. E. Parry, M. Reding, J. J. Royall, J. Schulkin, P. A. Sequeira, C. R. Slaughterbeck, S. C. Smith, A. J. Sodt, S. M. Sunkin, B. E. Swanson, M. P. Vawter, D. Williams, P. Wohnoutka, H. R. Zielke, D. H. Geschwind, P. R. Hof, S. M. Smith, C. Koch, S. G. N. Grant, A. R. Jones, An anatomically comprehensive atlas of the adult human brain transcriptome. Nature 489, 391–399 (2012).

11. B. A. Seitzman, C. Gratton, S. Marek, R. V. Raut, N. U. F. Dosenbach, B. L. Schlaggar, S. E. Petersen, D. J. Greene, A set of functionally-defined brain regions with improved representation of the subcortex and cerebellum. NeuroImage 206, 116290 (2020).

12. A. Arnatkevičiūtė, B. D. Fulcher, A. Fornito, A practical guide to linking brain-wide gene expression and neuroimaging data. NeuroImage 189, 353–367 (2019).

13. J. Quackenbush, Microarray data normalization and transformation. Nat Genet 32 **Suppl**, 496–501 (2002).

14. M. Hawrylycz, J. A. Miller, V. Menon, D. Feng, T. Dolbeare, A. L. Guillozet-Bongaarts, A. G. Jegga, B. J. Aronow, C.-K. Lee, A. Bernard, M. F. Glasser, D. L. Dierker, J. Menche, A. Szafer, F. Collman, P. Grange, K. A. Berman, S. Mihalas, Z. Yao, L. Stewart, A.-L. Barabási, J. Schulkin, J. Phillips, L. Ng, C. Dang, D. R. Haynor, A. Jones, D. C. Van Essen, C. Koch, E. Lein, Canonical genetic signatures of the adult human brain. Nat Neurosci 18, 1832–1844 (2015).

15. B. B. Avants, N. J. Tustison, G. Song, P. A. Cook, A. Klein, J. C. Gee, A reproducible evaluation of ANTs similarity metric performance in brain image registration. Neuroimage 54, 2033–2044 (2011).

16. G. Gryglewski, R. Seiger, G. M. James, G. M. Godbersen, A. Komorowski, J. Unterholzner, P. Michenthaler, A. Hahn, W. Wadsak, M. Mitterhauser, S. Kasper, R. Lanzenberger, Spatial analysis and high resolution mapping of the human whole-brain transcriptome for integrative analysis in neuroimaging. Neuroimage 176, 259–267 (2018).

17. E. T. Rolls, C.-C. Huang, C.-P. Lin, J. Feng, M. Joliot, Automated anatomical labelling atlas 3. Neuroimage 206, 116189 (2020).

18. B. D. Fulcher, M. A. Little, N. S. Jones, Highly comparative time-series analysis: the empirical structure of time series and their methods. Journal of The Royal Society Interface 10, 20130048 (2013).

